# Pseudodynamics+: Reconstructing Population Dynamics from Time-Resolved Single Cell Landscapes with Physics Informed Neural Networks

**DOI:** 10.64898/2025.11.30.691399

**Authors:** Weizhong Zheng, Melania Barile, Nicola K. Wilson, Yuanhua Huang, Fabian J. Theis, Berthold Göttgens

## Abstract

Single-cell profiling provides snapshots of the heterogeneous states that characterise developmental processes, organ regeneration and progression towards disease in a complex landscape. The underlying trajectories are of pivotal interest, but existing methods for reconstructing cell state trajectories commonly neglect population sizes. However, snapshot experiments make it difficult to interpret cell flux because the observed trajectories are confounded by changes in overall population size. This ambiguity can lead to misinterpreting changes in proliferation or death rates as changes in cellular migration. We introduce *pseudodynamics+*, a physics-informed neural network framework that solves high-dimensional flow equations on complex, branching landscapes. By integrating single-cell genomics with population dynamics, pseudodynamics+ estimates state- and time-dependent parameters of growth, differentiation, and diffusion. The model recapitulates proliferation bursts during T-cell maturation and, when applied to LARRY-barcoded data, predicts differentiation rates consistent with clonal behaviour. When applied to time-resolved persistent-labelling datasets of in vivo mouse bone marrow haematopoiesis, pseudodynamics+ reconstructs continuous tissue flows with dynamic parameters aligned with known molecular signatures. Notably, simulations revealed a previously unrecognised shift from megakaryocyte-biased to balanced progenitor output, explained by evolving fate preferences of progenitor states, as revealed by simulations leveraging our estimated dynamic parameters. Pseudodynamics+ therefore establishes a population-aware framework for reconstructing single-cell population dynamics and is available at https://github.com/Gottgens-lab/pseudodynamics_plus.

## Introduction

Time-series single-cell sequencing has enabled detailed characterization of developmental and cellular differentiation landscapes, providing deep molecular profiles to finely resolve cell states. If performed at sufficient scale, such time-series experiments should help to transform our understanding of tissue or population scale dynamics, by bridging the gap between single cell omics and conventional flux modelling, which has traditionally been agnostic of the deep molecular profiling information. However, precisely defining tissue-scale dynamic behaviour from single cell omics remains challenging. Firstly, the characterized cell states represent a discretisation of the continuous developmental process, whereby the destructive nature of current sequencing methods results in a loss of trajectory information important for connecting the transiting cell states. Secondly, despite the efficiency of single-cell experiments in exploring cell state space, captured cells may not accurately reflect actual cell numbers in the tissue, especially in rapidly growing systems (McDole et al., 2018; Fischer et al., 2019; Mittnenzweig et al., 2021). Consequently, single-cell snapshots may lose information about the exact expansion and shrinkage of the entire population. Moreover, both of these challenges are compounded when attempting to define complex multi-lineage differentiation landscapes.

Computational methods employing various principles have been proposed to connect cells across multiple time points. Optimal transport provides a sound mathematical foundation for learning ancestor-descendant relationships, such as Waddington-OT (Schiebinger et al., 2019) and moscot Klein et al. (2025b), by optimising the most energy-efficient cell-cell coupling between two single-cell datasets (Schiebinger et al., 2019; Bunne et al., 2023, 2024; Alatkar and Wang, 2023). While the optimised coupling can indicate transition paths by linking cells from two consecutive time points, it remains a static description. CellRank2 (Weiler et al., 2024) allows interrogation of cell state ordering using pseudotime to fill in the intermediate steps of the path. Alternatively, cellular trajectories can be calculated by generative models that predict the next state based on the current one. These models can generate transiting cell states, including those unobserved at intermediate time intervals, potentially imputing a fully continuous cell state trajectory. Several methods have been proposed based on this approach (Yeo et al., 2021; Chen et al., 2018a; Lipman et al., 2022; Tong et al., 2023; Klein et al., 2025a). Lastly, dynamic optimal transport-based methods have emerged as an advanced and continuous way to model single-cell distribution shifts across time points (Tong et al., 2020; Huguet et al., 2022; Sha et al., 2024). By solving the dynamic equation describing cellular density changes, methods adopting this concept not only learn the velocity field of continuous cell state transitions but also estimate the growth rate of individual cells. Nevertheless, none of the models mentioned above account for total population sizes, which can lead to dynamic parameters that do not reflect real-world physiological values.

Population dynamics, conversely, focus on modelling the temporal change of the exact total population size, providing an accurate quantification of tissue expansion and shrinkage. Computational modelling of population dynamics has traditionally been performed without access to single cell omics data (Frankovits et al., 2024), for example, by utilising flow cytometry analysis (Busch et al., 2015). Recently, however, *pseudodynamics* (referred to as *pseudodynamics-v1* later) was introduced as a mathematical framework that integrates population size measurements with single-cell omics snapshot data to infer tissue-level dynamics (Fischer et al., 2019). In its current formulation, *pseudodynamics-v1* reduces the continuous gene expression space to pseudotime coordinates— a discretization necessary for solving the governing partial differential equation (PDE). This simplification, however, limits the framework’ s capacity to resolve complex multi-branch systems, as cells occupying identical pseudotime positions across divergent developmental trajectories remain indistinguishable. Recent approaches address this limitation by modelling dynamic behaviour in multi-branching systems through cell-type or cluster-level counts scaled according to tissue size observations (Kucinski et al., 2024; Gao et al., 2024). Such methods reduce system complexity by representing cellular states as a finite set of discrete compartments, which can be described by ordinary differential equations (ODEs) that are computationally tractable. Nevertheless, tissue-scale dynamic modelling remains elusive at single-cell resolution due to the inherent challenges of solving PDEs for high-dimensional single-cell data (Zhu et al., 2024).

To address these limitations, we introduce *pseudodynamics+*, a computational framework that bridges time-series single-cell data and tissue size measurements to reconstruct complex population dynamics in a high-dimensional state space. Central to *pseudodynamics+* is a physics-informed neural network (Raissi et al., 2019; Cuomo et al., 2022) architecture that solves the governing partial differential equation (PDE) system without discretization, enabling joint estimation of time- and state-resolved parameters, including proliferation, differentiation, and diffusion rates at single-cell resolution. When applied to thymocyte maturation data, *pseudodynamics+* outperformed the original *pseudodynamics-v1* by resolving growth and differentiation rates that aligned better with both quantitative T-cell population trends (e.g., proliferative bursts during beta-selection) and independent biological observations. Benchmarking against state-of-the-art generative models using a LARRY-barcoded in vitro differentiation dataset demonstrated that *pseudodynamics+* achieves comparable accuracy to flow-matching approaches in predicting cell fate and generating future cell states. Finally, we applied *pseudodynamics+* to a uniquely complex in vivo dataset tracking the multi-lineage progeny of mouse haematopoietic stem cells for 9 months by combining transgenic label propagation with single-cell profiling. *Pseudodynamics+* enabled us to quantitatively track a multi-lineage continuum landscape in real-time, as well as inferring progenitor cell output bias via dedicated continuous density transport analysis, uniquely enabled by the dynamic behaviour characteristics revealed by *pseudodynamics+*.

## Results

### *pseudodynamics+* construct continuous tissue flow from single-cell time-series data

In developing biological systems, the distribution of cells along developmental trajectories evolves temporally, concurrent with tissue-scale population expansion or contraction (Fig. 1a). This dynamic process is captured by a time-dependent density function *u*(*s, t*), where *s* ∈ *S* denotes the cell state (e.g., low-dimensional transcriptomic coordinates) and *t*= 1,2, *· · ·, T* corresponds to experimental time points. Critically, we define *u*(*s, t*) as an unnormalized density whereby its integral over *S* recapitulates the total population size *N*_*t*_ = ⎰_*S*_ *u*(*s, t*)*ds*. The total number of cells in the population, which should be measured additionally,. This formulation bridges single-cell state transitions inferred from sequencing data with tissue-level dynamics, enabling quantitative modelling of proliferation, differentiation, and stochastic state dispersion. To reconstruct single-cell-resolved dynamics, we propose *pseudodynamics+*, a computational framework modelling cellular density governed by a partial differential equation (PDE).

**Figure 1.**
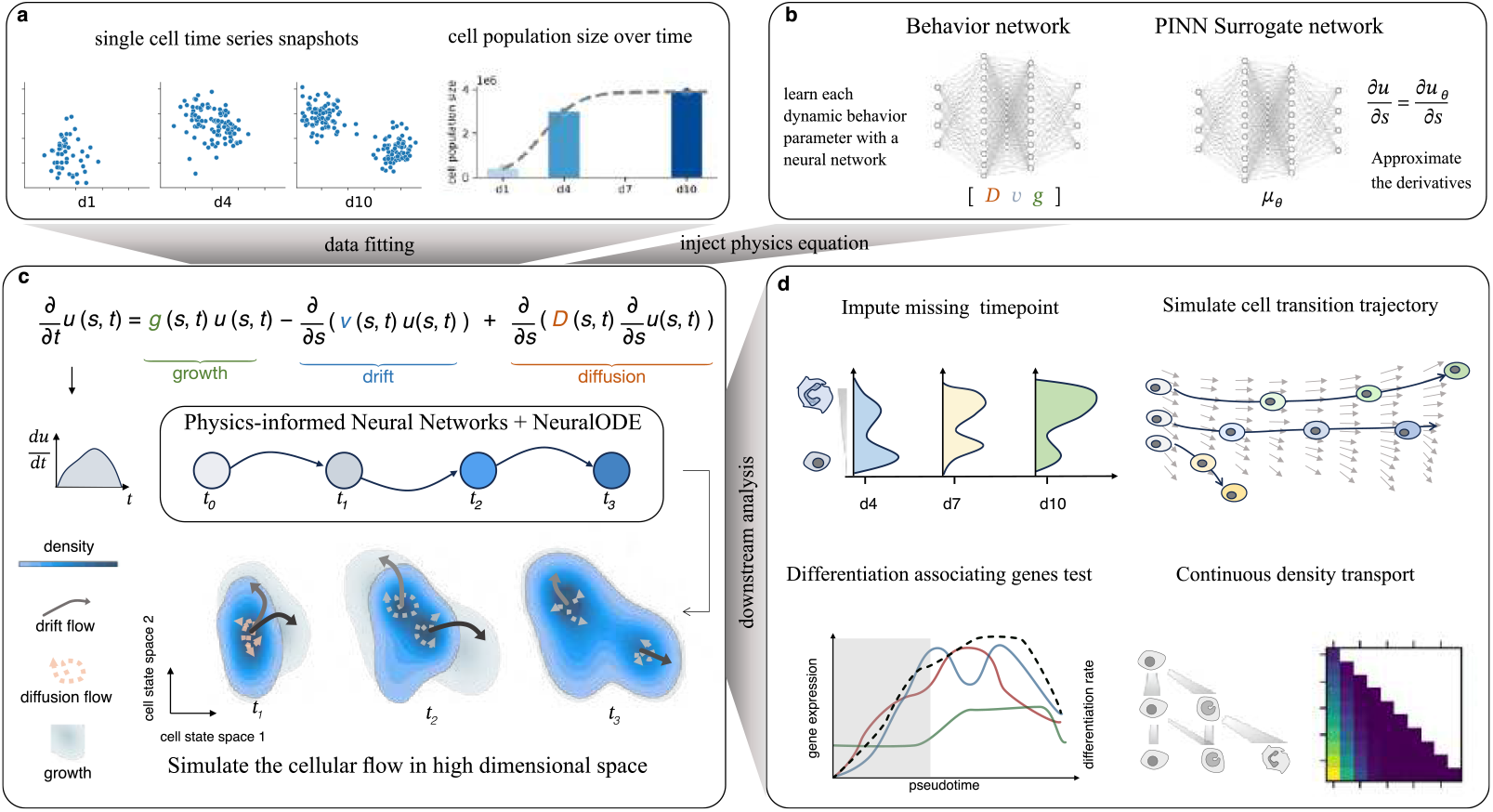
Overview of the pseudodynamics+ Model Framework. **(a)** *pseudodynamics+* takes single-cell time-series data and corresponding population size as input. **(b)** The model utilizes surrogate and behaviour neural networks to parameterize density and dynamic rates (growth, differentiation, diffusion), respectively. **(c)** A Neural ODE solver integrates these components to solve the governing partial differential equation. **(d)** Downstream analyses enabled by *pseudodynamics+*, including timepoint imputation, cell state trajectory simulation, differentiation associated gene test and continuous density transport.”

The temporal change in density, 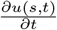, in this advection-reaction-diffusion equation is determined by three dynamic behaviours: growth, drift, and diffusion (Fig. 1b). Together, these behaviours characterise how a cell, with state s at time t, will evolve in the next moment. These behaviours are controlled by three parameters, respectively: 1) the net proliferation rate, *g*(*s, t*), resulting from cell division and cell death; 2) the velocity, *v*(*s, t*), defines the speed and direction of deterministic cell state transition due to differentiation; and 3) the diffusion coefficient, *D*(*s, t*), controls the degree of stochastic density dispersion among neighbouring cell states. By taking single-cell sequencing data and tissue size as input, *pseudodynamics+* infers the time- and cell state-specific dynamic parameters for each cell, with which we can reconstruct the continuous cell flux underlying the developing tissue.

Solving such a PDE for single-cell data is non-trivial, as it requires both the first- and second-order derivatives of density *u* with respect to the high-dimensional cell state *s*. To address this problem, *pseudodynamics+* approximates the density function using Physics-Informed Neural Networks (PINNs), *u*_*θ*_ (the surrogate network), a method that uses neural networks to solve mesh-free high-dimensional PDE systems (Raissi et al., 2017, 2019; Haghighat et al., 2021; Torres et al., 2022). Leveraging this technique, we can estimate the derivative terms 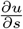 and 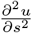 obeying Equation 1. Moreover, we parameterize each of the behaviour functions *g, v*, and *D* with a neural network (the behaviour network) dependent on *s* and *t*. Consequently, the density dynamics (the left-hand side of the PDE Equation 1) can be reformulated as the combination of the output from four neural network modules, which makes it feasible to solve using neural ODEs (see Method, Fig. 1c).

The inferred state- and time-resolved parameters enable quantitative analysis of the dynamic processes (Fig. 1d). A direct application is to impute the unobserved intermediate time points, in terms of the density distributions *u*(*s, t*) and total tissue population sizes *N*_*t*_. The drift term *v*(*s, t*) encodes the velocity field of directed cell state transitions, which — when combined with the differentiation rate,— allows simulation of individual cell trajectories and temporal outflux from specific states. By integrating the growth rate *g*(*s, t*) with differentiation dynamics, we compute ancestor-descendant transition probabilities across the trajectory, generating a continuous density transfer map. This map quantifies redistribution of cellular densities between consecutive time points, revealing how lineage-specific expansion, contraction, or differentiation drives the temporal evolution of population-level dynamics.

For simplicity, the cell state *s* used as input is a low-dimensional representation that effectively retains the topology of cell connectivity. This is based on the assumption that the gene expression of cells sequenced in single-cell experiments lies on a low-dimensional manifold.

### Benchmarking density estimation on single-lineage data and evaluating parameter estimation on synthetic data

The *Pseudodynamics+* framework models cellular differentiation as a dynamic system, where a continuous cell mass drifts across a developmental landscape over time. Accurately quantifying this cellular density from high-dimensional single-cell data is non-trivial. To obtain a precise density estimate for fitting a Physics-Informed Neural Network (PINN) at observed time points, we first benchmarked various density estimation methods and low-dimensional representations for their ability to capture the progression of cell mass along a differentiation trajectory. We used a megakaryocyte differentiation dataset Kucinski et al. (2024) for this evaluation, as it profiles a clear transition from hematopoietic stem cells (HSCs) to progenitors and finally to megakaryocytes (Fig. S. 1a), with a prominent shift in cell states between Day 3 and Day 49 (Fig. S. 1b).

For this single-lineage dataset, we defined the density estimated along a one-dimensional pseudotime axis (as used in *pseudodynamics-v1*) as the ground truth. We compared several density estimation methods, including: traditional Gaussian Kernel Density Estimation (KDE), hashing-based KDE Charikar and Siminelakis (2017), the Gaussian Mixture Model from TIGON Sha et al. (2024), the deep learning-based method Denmarf Papamakarios et al. (2017), and the recently proposed Gaussian Process-based method Mellon Otto et al. (2024). These methods were evaluated on both diffusion map (DM) and principal component (PC) spaces, with the resulting densities subsequently projected back to the pseudotime axis for comparison. Overall, the results indicate that density estimates derived from diffusion map coordinates achieve higher concordance with the pseudotime-based ground truth. Although TIGON’ s estimator achieved a high correlation score, it failed to accurately recover the density accumulation in stem cells on Day 3 and the subsequent mass transition to progenitor states on Day 7 (Fig. S. 1c-d). Consequently, we selected diffusion map coordinates coupled with traditional Gaussian KDE, as this combination yielded cell densities that best captured the distinct developmental stages and the temporal progression of dense regions (Fig. S. 1c-f).

Next, we assessed the accuracy of *Pseudodynamics+* in inferring dynamic parameters using synthetic five-dimensional time-series data with known ground-truth parameters. We first generated ground-truth velocity functions by fitting five natural cubic splines with randomly sampled knots, one for each data dimension. The ground-truth growth and diffusion rate functions were defined with other sets of spine functions but connected with linear combinations, resulting in scalar values that change over time. We then simulated cell states by propagating points according to the velocity functions and computed the resulting density by performing an integral over the underlying partial differential equation (PDE) system, incorporating all three dynamic parameters. Finally, we fitted the *Pseudodynamics+* model to this synthetic data and compared the PINN-inferred parameters to the ground truth. As shown in Fig. S. 2, *Pseudodynamics+* accurately estimated the growth rate, recovering its increasing trend over time. For the differentiation rate, we compared the estimated and ground-truth values for each dimension, achieving an average Pearson correlation of 0.81 (Fig. S. 2d).

### *pseudodynamics+* identified multiple waves of proliferative burst of T-cell maturation

We applied our method to study the dynamic process of T-cell maturation in the mouse embryonic thymus using a previously published single-cell RNA sequencing dataset containing eight time points evenly sampled from embryonic day (E) 12.5 to E19.5 in the mouse embryonic thymus (Kernfeld et al., 2018). This single-cell data captured around 48,000 cells, tracing the T-cell trajectory from progenitor cells, double-negative (DN) to more mature double-positive (DP) cells, as well as a small branch of non-canonical lymphoid (NCL) cells (Fig. 2b). The total population size of each embryonic stage was counted separately (Cook, 2010). When the same dataset was used in the original *pseudodynamics-v1* paper (Fischer et al., 2019), cells were ordered along the pseudotime coordinate, and cell states were defined as pseudotime bins to model the continuous density transition process of the T-cell lineage (Fig. 2ac). However, this simplification means that pseudotime bins might be contaminated with cells from different stages or terminally differentiated cells (e.g., Phase 2-2 cells) that would not progress to downstream progeny. Consequently, the parameters estimated by *pseudodynamics-v1* would reflect the aggregated properties of functionally distinct cells. In contrast, we trained our *pseudodynamics+* in a multi-dimensional diffusion map space, enabling us to learn the dynamic behaviour of each single cell within a complex multi-lineage differentiation manifold.

**Figure 2.**
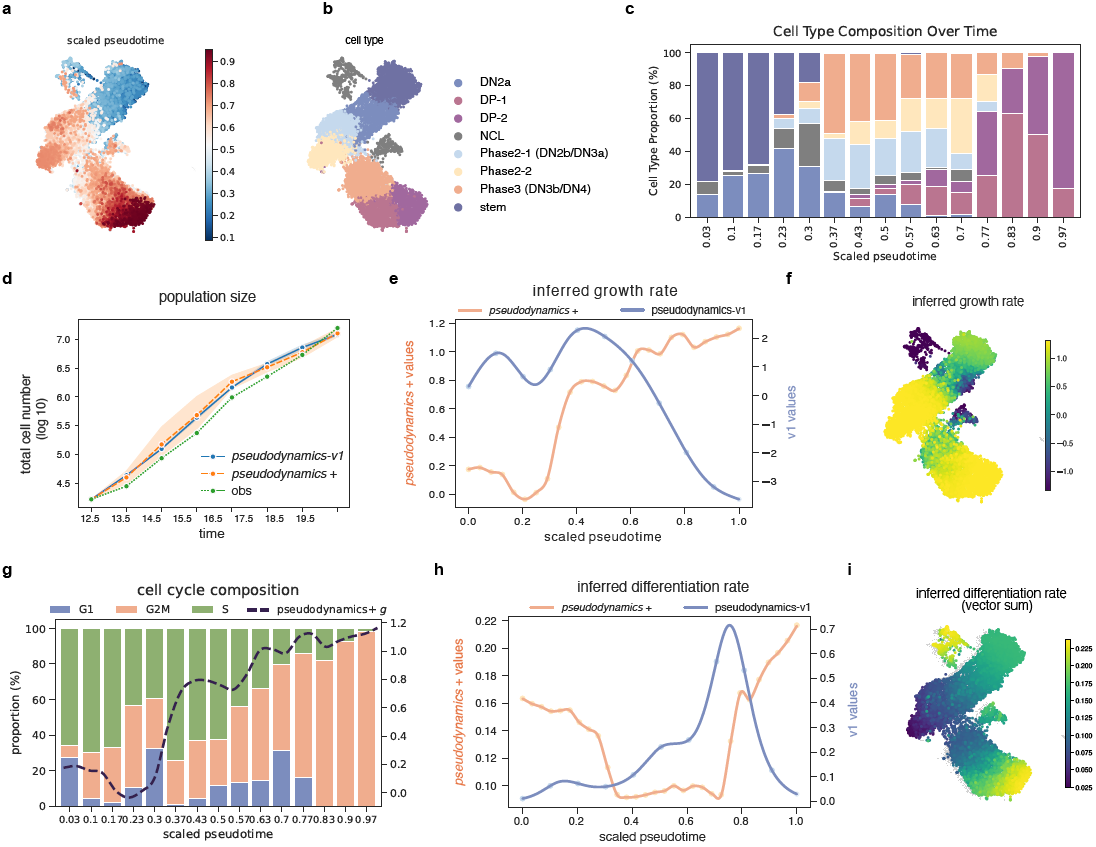
Dynamic Parameter Inference in Thymus Development. **(a, b)** UMAP embeddings of single-cell data, illustrating scaled pseudotime and cell type annotations. **(c)** Cell type composition across pseudotime bins, with the same colour regime as **(b)**. **(d)** Total cell number (log10) over time, comparing observed data with predictions from *pseudodynamics-v1* and *pseudodynamics+* models. **(e, f)** Growth rate profiles plotted on UMAP space (e) and along pseudotime (f). **(g)** Distribution of cell cycle phases across pseudotime intervals. **(h, i)** Inferred differentiation rate profiles along pseudotime and projected onto UMAP space. A multi-dimensional drift vector is normalized within each dimension and then summed across dimension to turn into a scaler. DN – double negative, DP – double positive, NCL - non-canonical lymphoid

Immature T-cells in the embryonic thymus undergo active expansion, driven by several rounds of selection that induce extensive programmed cell death of unqualified cells (XIAO et al., 2003; Park et al., 2020b). Examining the estimated total population size, both versions of *pseudodynamics* can recover the rapid expansion of thymocytes at the tissue scale (Fig. 2d). Visualising Pseudodynamics+ growth rates in UMAP space (Fig. 2f) showed reduced proliferation of phase-transiting cells around the DN2a and Phase-2/3 transitions, indicating diminished surface-protein selection of T-cells. These low-proliferative cell states aligned with the pseudodynamics-v1-inferred growth rates, which exhibited two minima around pseudotime 0.2 to 0.6 (Fig. 2e). Pseudodynamics+ also identified three waves of proliferative bursts, namely in the progenitor, Phase 2, and DP cells. In contrast, pseudodynamics-v1 only suggested the first two waves and assigned a decreasing growth rate as cells progressed towards the DP phase. However, cell cycle phase prediction based on scRNA-seq analysis showed a high G2M proportion of cycling DP cells (Fig. 2g), thus validating the third wave of proliferative burst predicted by Pseudodynamics+.

The vectorial sum of the inferred differentiation rate provides each cell a single value denoting its instantaneous change in multi-dimensional diffusion map coordinates. *Pseudodynamics+* revealed rapid cell state transitions occurring both near the root and the terminus of the T-cell trajectory (Fig. 2h). Notably, the lowest differentiation rate was assigned to the tip of the Phase 2-2 cells, which was largely disconnected from downstream cell identities, likely representing a differentiation dead end that failed to pass *β*-selection (Fig. 2i). Compared with pseudodynamics-v1, our aggregated differentiation rate demonstrated a distinct pattern, particularly at the end of the trajectory (Fig. 2h). Whereas pseudodynamics-v1 enforced cells to stop differentiating at the boundary, our method predicted a rapid differentiation rate and allowed cells to transition to even further downstream cell states. This difference arises from our assumption that the system is not closed, in which there exists outflow from DP cells to cell states that are not included in the sequencing data. Indeed, it is well-established that DP cells will further differentiate and undergo another round of selection to become single positive T-cells (Leavy, 2007; Ross et al., 2014).

Pseudodynamics+ is uniquely powered to make predictions about future cell densities. Across different time points, *pseudodynamics+* accurately simulated the density distribution based on that of the previous time point (Fig. S. 3). The learned density transition over time reconstructed the complex dynamic behaviour of the system. This reconstruction began with the early expansion of progenitor cells and differentiation into DN2a-phase cells from E12.5 to E14.5. Subsequently, the reconstructed density showed the bifurcation of Phase 2-2 cells at E15.5, with an accumulation of cells before and after *β*-selection. From E16.5 to E19.5, *pseudodynamics+* predicted the accumulation of DP-phase cells with a gradual transition in the cell state space (Fig. S. 3c). These results indicate that the dynamic parameters learned by *pseudodynamics+* can accurately describe the profiled T-cell development system in this dataset, capturing key aspects such as branching that cannot be accounted for by the original method. In summary, *pseudodynamics+* learned a set of dynamic parameters that not only explained the fast-growing population and density changes during thymocyte maturation but also aligned with the underlying biology in regard to the cell cycle composition and the possible transition to downstream maturation stages.

### Velocity field inference by *pseudodynamics+* aligns with lineage tracing data

As outlined above, *pseudodynamics+* allows us to learn the velocity field of the cell representation space. The resulting behavior network *ν*_**w**_ can therefore act like a continuous-flow model, which can be used to simulate the nature of future cell states given the starting cell. We next wanted to explore how well *pseudodynamics+* performed when analysing time-resolved differentiation landscapes that had incorporated ground truth on actual differentiation paths taken by individual cells. To this end, we turned to the dataset reported in a landmark study (Weinreb et al., 2020) tracking in vitro differentiation of mouse haematopoietic stem/progenitor cells using the LARRY molecular barcodes. LARRY barcodes are stably introduced into cells at the start of the time course experiment and can be read out by scRNA-Seq, thus provide a direct means to assign cellular ancestries. For comparison with existing tools, *pseudodynamics+* was benchmarked against a range of tools, including the recurrent neural network based PRESCIENT, dynamic optimal transport based methods TrajectoryNet, MIOFlow and TIGON, and the latest flow-matching based model OT-cfm and SF2M. All the models were trained within the 5 dimensional diffusion map space.

We first evaluated the accuracy of fate bias prediction across methods, assessing clonal membership of cells across the three measured timepoints at days 2, 4 and 6. For cells of the same clone, we computed the cell type distribution of the clone at the last time point (Day 6) and then defined this distribution to cells at the first time point (Day 2) as their fate bias. Using the labelled initial cells from Day 2, we used the previously mentioned tools to simulate their cell states at Day 6. Cell type annotations of these simulated cells were transferred from the nearest cells in the actual dataset. As shown in Fig. 3a, *pseudodynamics+* outperformed most existing methods and achieved comparable accuracy with the latest flow-matching models. Interestingly, dynamic optimal transport-based methods underperformed, possibly due to their primary goal of reconstructing density changes rather than predicting cell state.

**Figure 3.**
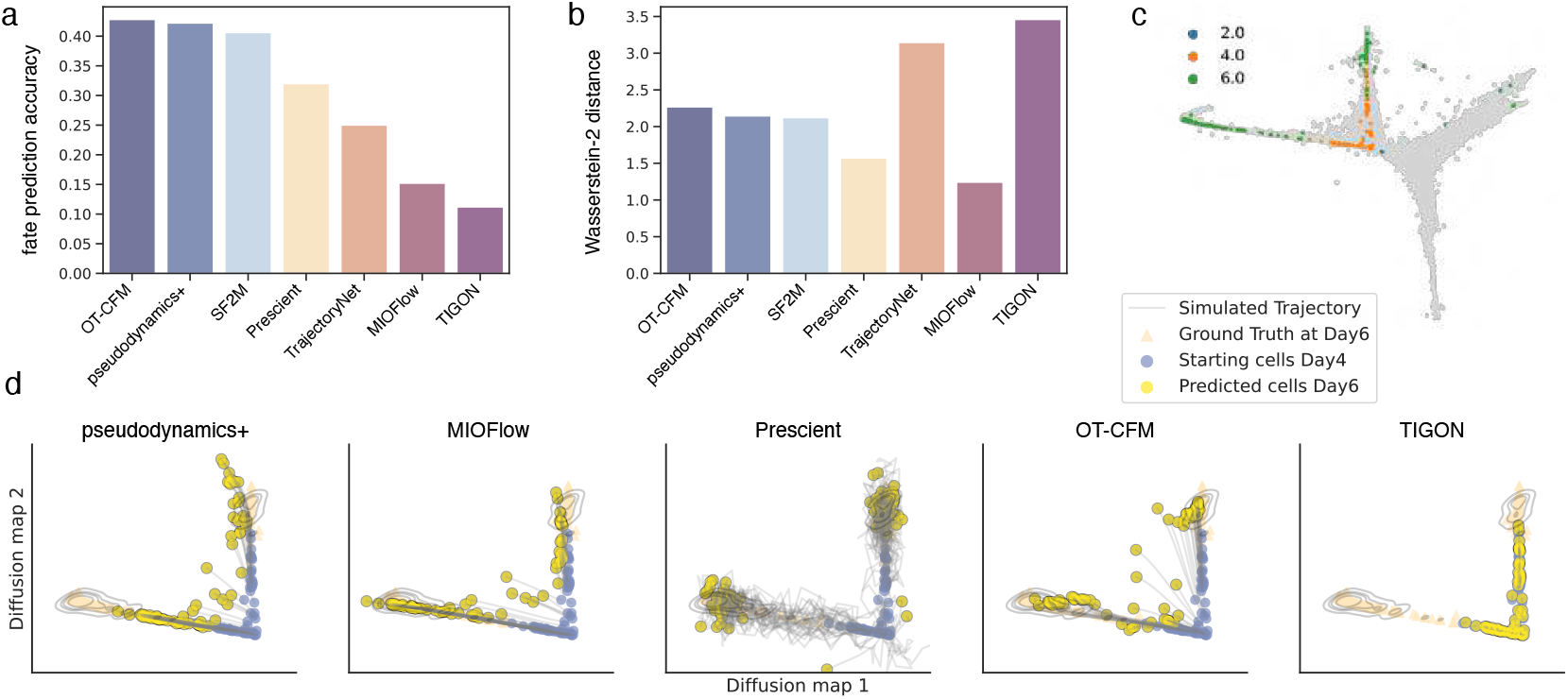
Benchmarking Cell Trajectory Prediction Using LARRY Barcoded cultured Haematopoietic Cells. **(a)** Accuracy of cell fate prediction across different methods. Higher values indicate better performance. **(b)** Wasserstein distance (W2) between simulated and observed cell state coordinates at Day 6, reflecting trajectory accuracy. Lower values indicate better performance **(c)** Visualization of selected clones in the first two diffusion map dimensions, used for trajectory analysis. **(d)** Simulated cell trajectories from Day 4 to Day 6 for individual methods. Blue points: starting cell positions at Day 4, Yellow points: predicted cell positions at Day 6, Orange contours: observed cell density at Day 6.

Cell fate prediction, as performed above, evaluates the simulation endpoints based on the assignment of cell type categories—a rather coarse-grained measurement given that we are dealing with continuous differentiation landscapes. To quantify the similarity between simulated and ground-truth cell populations in a more high-dimensional space, we calculated the Wasserstein distance for cells from the same clone. We selected the two largest clones with cells falling into multiple branches, specifically the monocyte and neutrophil lineages (Fig. 3c). In this task, *pseudodynamics+* performed similarly to the flow-matching methods; yet, it was surpassed by Prescient and MIOFlow. In the diffusion map space, *pseudodynamics+* showed the correct direction of differentiation. Additionally, in both our method and OT-CFM methods, the simulated trajectories contained intermediate branching events, effectively reconstructing the highly continuous differentiation processes and aligning with the overall distribution. In comparison, Prescient produced highly accurate yet inherently stochastic trajectories. However, several cells diverged from the main branches and accumulated in undefined regions of the diffusion map.

In summary, the velocity field learned by *pseudodynamics+* can yield accurate cell state trajectories and demonstrates highly competitive precision compared to state-of-the-art flow-matching methods.

### Single cell resolved flux model for long-term mouse in vivo haematopoiesis

Characterising the dynamics of in vivo haematopoiesis is challenging, particularly in the context of steady-state unperturbed homeostasis (Busch et al., 2015). Despite the substantial daily turnover of blood cells, the distribution of different blood cell compartments is tightly regulated to remain stable. This regulation makes it difficult to quantify the number of newly synthesized cells and the time required for immature progenitor cells to differentiate. To address this problem, we used a recently published single-cell sequencing dataset that employs persistent labelling. This method could inducibly and permanently label haematopoietic stem cells (HSCs) and all of their subsequent progenies via expression of a red fluorescent tdTomato transgene. Fluorescent cells from the bone marrow of label-induced mice were then profiled by scRNA-Seq, spanning a broad timespan from 3 days to 269 days post-induction, allowing us to measure the in-flux and out-flux across the refined haematopoietic landscape across time to be measured. The dataset captured a time-resolved differentiation landscape made up of HSCs, intermediate multi-lineage progenitors, and lineage-committed progenitors (Kucinski et al., 2024) (Fig. 4a). Applying *pseudodynamics+*, we then inferred parameters capturing the dynamic behaviour of the in vivo mouse haematopoietic stem and progenitor compartment.

**Figure 4.**
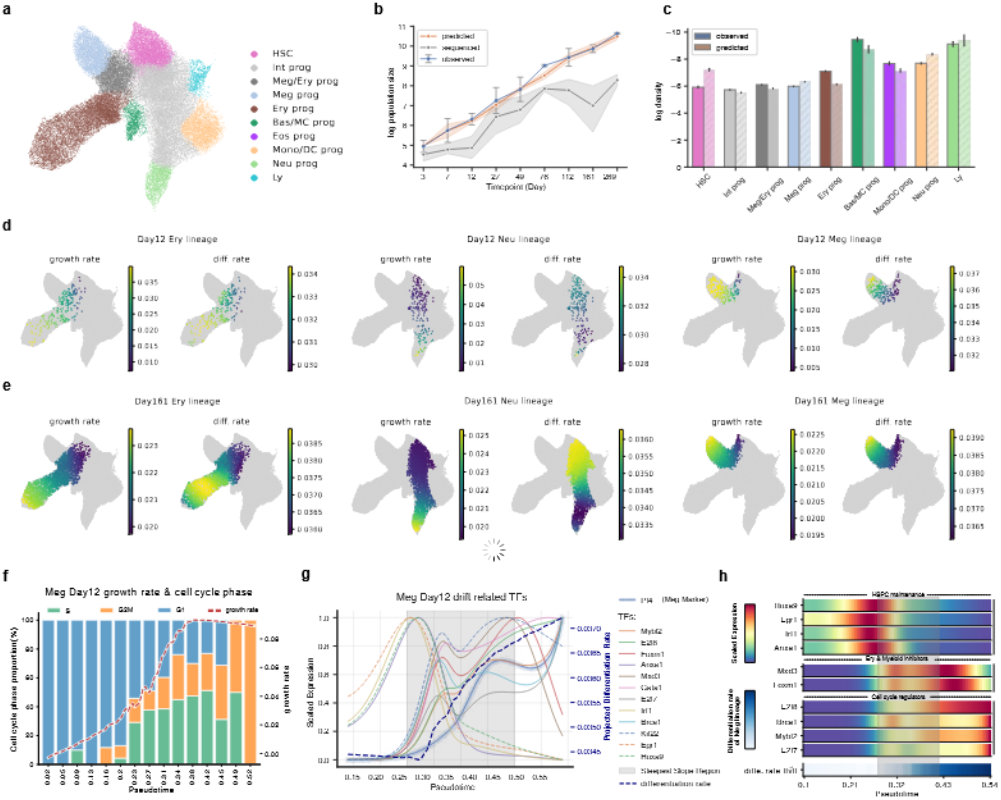
Analysis of dynamic parameters in persistent labelling data of in vivo mouse haematopoiesis. **(a)** UMAP visualisation illustrating cell type annotations. **(b)** Line plot depicting the total cell number (log-transformed) as a function of time. The blue curve represents the observed population size, with error bars indicating the standard deviation (std) derived from experimental replicates. The orange curve shows the simulation results obtained using *pseudodynamics+*, with error bars representing the standard deviation across multiple training repeats. The grey curve illustrates the observed number of sequenced cells. **(c)** Bar plot evaluating the imputed cell density, aggregated by cell type at Day 161. The colour scheme is consistent with (a), and solid and hashed patterns denote observed and predicted densities, respectively. **(d, e)** UMAP visualisations displaying the estimated growth and differentiation rates for megakaryocyte (meg), erythroid (ery), and neutrophil (neu) lineages at Day 12 and Day 161. Lineage annotation of the cells is assigned based on their fate probability, as determined by (Kucinski et al., 2024). **(f)** Overlay of projected growth rate of Meg lineage at Day 12 (dashed red curve) and cell cycle phase composition of across pseudotime. **(g)** The fitted gene expression curve as a function of pseudotime. In the plot we show top 12 differentially expressed TFs that are ranked by the absolute log fold change. The bold dashed line represent the projected differentiation rate for Meg lineage at Day12 and the shaded region stands for the pseudotime interval with the steepest slope. **(h)** Heatmap showing the expression pattern of three TF groups across the pseudotime. TFs are selected with the same association test as in (g). HSC – haematopoietic stem cell; Int prog – intermediate progenitor cells; meg prog megakaryocyte progenitors; Ery prog – erythrocyte progenitors; Bas/MC – Basophil/mast cell progenitors; Eos – eosinophil progenitors; mono/DC – monocytic/dendrocytic progenitors; Neu – neutrophil progenitors; Ly – Lymphoid progenitors.

Quantifying the total number of fluorescence-labelled cells across the landscape over time revealed a gradual slowdown in population expansion over time, indicating a previously unrecognised shift in the system’s dynamic properties (Fig. 4b). To model this trend, the static-rate setting adapted by previous studies (Fischer et al., 2019; Kucinski et al., 2024) needed to be extended to incorporate the modelling of time-sensitive dynamic parameters, which provided a good fit between the model and experimental data (average KLD = 0.136 for seen training timepoints, Fig. S. 5b,f). More importantly, the parameters estimated by *pseudodynamics+* were able to accurately imputed the cell-type density for two held-out timepoints, day 49 and 161 Fig. 4c, Fig. S. 5c-f, average KLD = 0.097). The two unseen timepoints represent an earlier transiting state when rates are changing, as well as a later more homeostatic state, thus demonstrating that our derived dynamic parameters are able to capture the long-term density change of the whole landscape across the entire 9 months’ timeframe.

Analysis of *pseudodynamics+*-estimated parameters revealed lineage-specific patterns of growth and differentiation (Fig. 4d). The Megakaryocyte and Erythroid lineage exhibited coupled growth and differentiation rates (Fig. 4d-e), both progressing from HSCs towards the terminal states. In contrast, the Neutrophil lineage demonstrated a a decoupled differentiation and growth rate pattern, with a decrease in differentiation from the intermediate progenitor state until the terminal point. Whilst the Erythroid parameters remained stable over the 9 months’ timeline (Fig. S. 6a), the Megakaryocyte lineage exhibited an early proliferative peak during Days 3–12, consistent with rapid replenishment. Growth rates and cell cycle phase prediction were closely aligned when projected onto pseudotime, an association observed across timepoints for both the Megakaryocyte and Erythroid lineages (Fig. 4f, Fig. S. 7a–b). Our time-sensitive *pseudodynamics+* model revealed a uniquely complex and evolving behaviour for the Neutrophil lineage. Early growth rates correlated well with cell cycle phase prediction, but this association was lost at later stages as the system transitioned into a homeostatic state (Fig. S. 7 ab), probably due to the shift in cell cycle activity.

To validate the estimated differentiation rate inferred by *pseudodynamics+*, we developed a pipeline to identify drift associating genes, with the assumption that meaningful parameters should correspond to known lineage markers. Basically, the pipeline performed differential expression test within pseudotime intervals of maximal differentiation rate change (the drift-variable state) and combined with correlation analysis along entire trajectories (Fig. S. 8a). This approach recovered canonical lineage regulators across all three lineages (Fig. S. 8b-g): Pf4 and Gata1 for Megakaryocytes (Lambert et al., 2014; Lee et al., 2017), Klf1 and Gata1 for Erythocytes (Gutiérrez et al., 2020; Lee et al., 2017; Siatecka and Bieker, 2011; Tallack and Perkins, 2010), and Cebpe and Cebpa for Neutrophils (Shyamsunder et al., 2018; Theilgaard-Mönch et al., 2022) (Fig. S. 8b–g). In addition, the association test also allows the identification of dynamic TF modules active at different stages of a given trajectory. In the Day 12 Meg lineage for example, we identified three groups of TFs peaking at different stages of the trajectory (Fig. 4h), implying divergent roles in promoting the maturation of megakaryocyte progenitor. Hoxa9, Egr1, Irf1, and Anxa1, linked to HSPC maintenance and myeloid differentiation (Ramos-Mejía et al., 2014; Rundberg Nilsson et al., 2023; Nilsson et al., 2023), were rapidly down-regulated following the early burst in differentiation rate. A second group of TFs which remained highly expressed for longer includes factors with known roles in cell cycle regulation (Di Stefano et al., 2003; Westendorp et al., 2012; Deng, 2006; Liu et al., 2023). The third group, including Foxm1 and Mxd3, known inhibitors of erythroid and myeloid differentiation respectively, showed transient peaks of expression before being silenced, suggesting a role in constraining Megakaryocyte fate by blocking alternative lineages. Taken together, Pseudodynamics+ inferred time-dependent dynamic parameters that not only recapitulated the density landscape but also aligned with lineage-specific molecular features, including cell cycle phase composition and the expression of lineage-defining genes.

### *pseudodynamics+* reveals the transition from quick megakaryocytic biased haematopoiesis to slow homeostatic haematopoiesis over time

We next characterized system-wide rate shifts across hematopoietic compartments over the nine-month time course. Firstly, we observed that relative differentiation rates remained stable after Day 76. Secondly, the megakaryocyte and erythrocyte lineages displayed rapid differentiation throughout the time course, while monocyte and dendritic progenitors exhibited progressively increasing differentiation rates (Fig. 5a). Of note, growth rates inferred for the early time points showed substantial variation, yet by day 76 had converged into a more stable value range (Fig. 5b). Meg, Ery, and Neu progenitor exhibited substantially elevated proliferation in early time points (Supp Fig. S. 7a), in line with measured behaviours seen in a previous study Upadhaya et al. (2018), where MK progenitors significantly outpaced other downstream progenitors in the first week post-tamoxifen. Importantly, our implementation of time dependent rate changes prevents any possible anomalies from the early time points corrupting the overall results. For example, *pseudodynamics+* predicts that HSCs divide rarely, whereas time-independent model assigned high growth rates to HSCs (Fig. S. 6c), incompatible with their known quiescence Takeishi et al. (2025) as well as the low cell cycle scores observed in single cell RNA-Seq.

**Figure 5.**
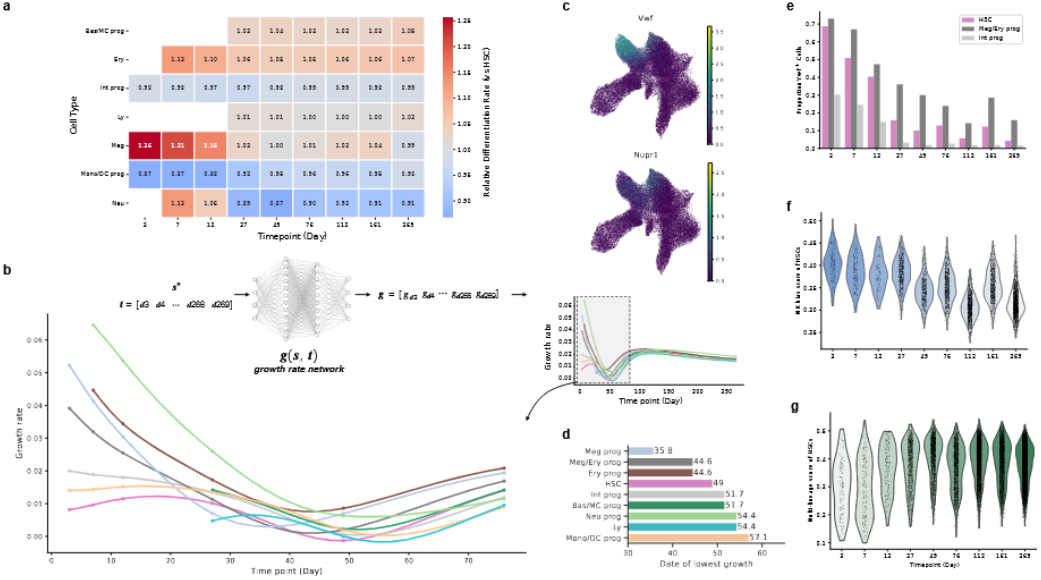
The temporal shift of lineage output revealed by the time-resolved dynamic parameters. **(a)**Heatmap showing the relative differentiation rate of each cell type compared with HSCs. Only cells from the cell the corresponding timepoint were used to aggregate and scaled over HSCs. Cell types with less than 10 sequenced cells were not shown in the heatmap. **(b)** Temporal profiles of growth parameters, aggregated by cell type over the experimental time course. The growth curve of individual cells between two collected timepoints is densely evaluated at 10 intermediate timepoints. Nodes on the curve indicate the observed timepoint **(c)** UMAP visualizations illustrating the expression levels of vWF and Nurp1. **(d)** Bar plot illustrating the timepoint (date) at which the minimal growth value occurs for each cell type. **(e)** Temporal distribution of vWF-expressing cells. **(f, g)** Violin plots representing the time-wise distribution of the megakaryocyte (MK)-biased gene signature score and the multi-lineage output score for HSCs.

Paralleling the early growth profile, a recently identified HSC subgroup restricted to produce megakaryocyte (vWF+ HSCs)was detected from Day 3 to Day 12 (Fig. 5c, f), Sanjuan-Pla et al. (2013); Carrelha et al. (2018); Haas et al. (2015), followed by a rise of the megakaryocyte marker Pf4 (Fig. 4g, Fig. 5c). Moreover, gene signature scoring also suggests a decreasing MK bias over time (Fig. 5f-g). To further explore the possible overrepresentation of MK-specialised HSPCs during the early time window, we interrogated our data with recently reported genes associated with the alternative Mk pathway, derived from single-cell expression comparisons of vWF+ HSCs with standard multi-lineage HSCs Carrelha et al. (2024), which showed significantly higher expression in our early-phase vWF+ HSCs (Fig. S. 9d, max p-value = 0.00735 using Mann–Whitney U test), an example of which is Nupr1 (Fig. 5c). We next processed the published data to identify the most differentially expressed genes between vWF+ and standard HSCs to define two corresponding gene sets. Our vWF+ HSC population, especially during the early phase, scored higher in the vWF+ P-HSC signature (Fig. S. 9b), consistent with a non-canonical trajectory towards megakaryocytes. By contrast, later-phase cells from Day 76 onwards displayed more prominent expression of the standard HSC signature genes (Fig. S. 9c). These results are consistent with a model whereby a subpopulation of MK-specialised HSCs has substantially faster transit times and therefore reaches immature progenitor stages first, thus appearing enriched in overall molecular profiles. Over time, most slower transiting cells catch up and therefore end up as the majority of cells throughout the landscape. Of note, none of these complex behaviours could have been revealed without allowing for time-dependent changes in flux parameters.

### Continuous density transport reveals dynamically distinct progenitors subgroups

To further explore how the heterogeneity within HSCs and the progenitor cells affect the dynamics of the landscape, we leveraged the learned dynamic parameters to develop a new framework, continuous density transport (CDT), allowing us to quantify how a cellular density redistribute stepwise among its progenies along the differentiation trajectory (Fig. 6a). Specifically, CDT firstly uses the learned velocity field *v* (*s, t*) to simulate the cell state transition trajectory [*s*_1_, *s*_2_, .. *s*_*n*_] over a time window *t*_1_ →*t*_*n*_ with several intermediate timepoint [*t*_1_, *t*_2_, … , *t*_*n*_]. For each time interval [*t*_*j*_, *t*_*j*+1_], we combined the drift term of the equation with the simulated cell state transition to estimate the density outflow 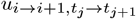 from a source cell *s*_*i*_ to its target state *s*_*i*+1_ (*i∈*[1. *n*]), together with the inflow 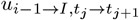 and density retention 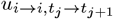. During simulation, fluxes are further modulated by the cell state–specific growth rate *g*(*s, t*). In this way, by combining several dynamic parameters at the same time, we compute each cell a transport map (Fig. 6a) which can delineate the cell flux between all of it progenies by its cell mass (log density in this study), the outset and the destination in tissue scale.

**Figure 6.**
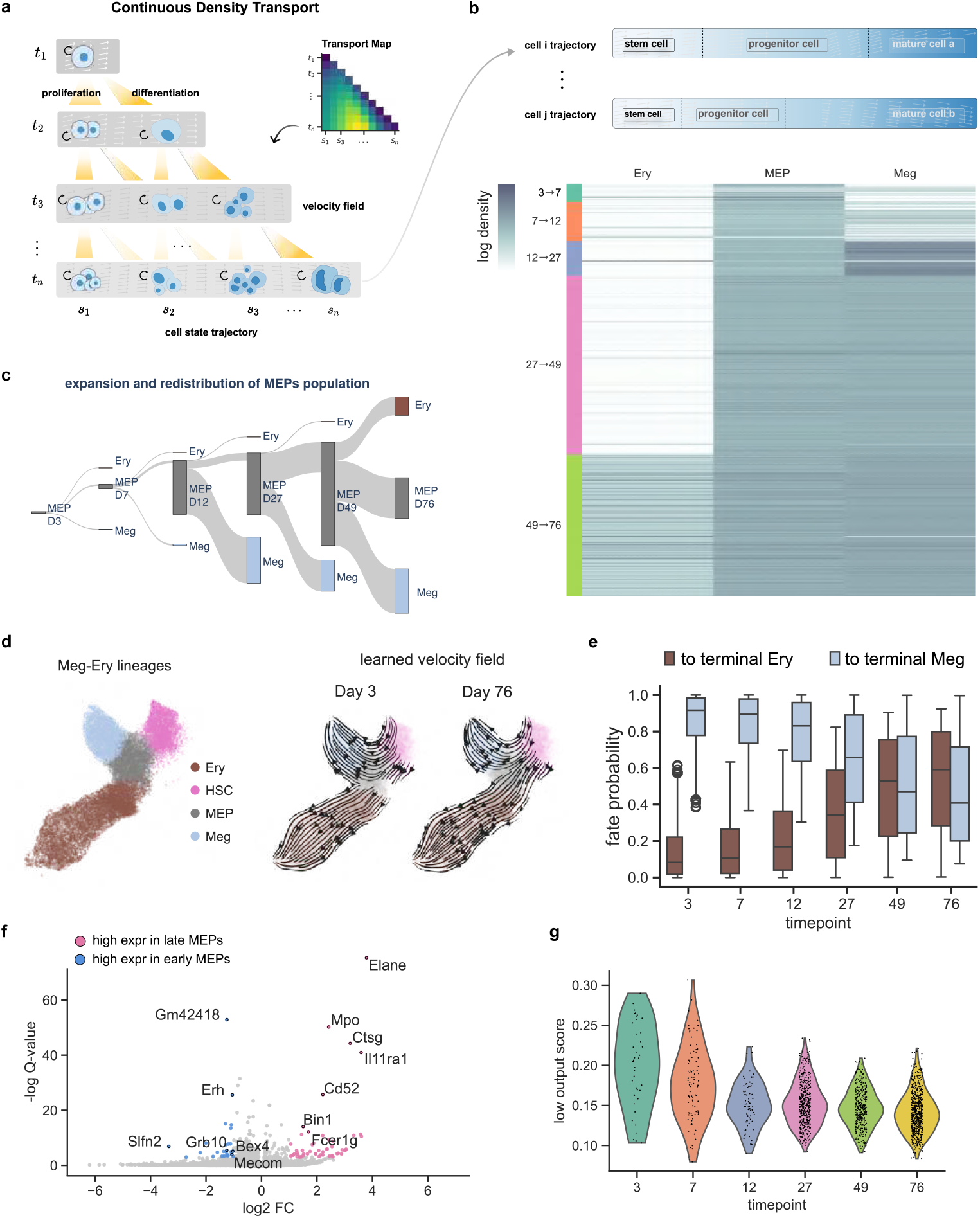
Characterization of HSC Functional Heterogeneity through Continuous Density Transport Analysis. **(a)** Schematic representation of continuous density transport analysis. For each cell, a simulated trajectory is generated, and the density flow along this trajectory is quantified. The x-axis represents the cell trajectory as it transitions to downstream cell states. From the top to the bottom, the time is divided into n intervals and the each row of the y-axis represents the time at which the density transport is evaluated. **(b)** Illustrative schema of cell type labelling for each simulated cell state and aggregation of density by cell types. Heatmap in the bottom shows the transition of density at the last step (*t*_*n*_ )from MEPs to other cell types according to the annotation. The density aggregates different cell state at different steps . Each row stands for a cell and the row sum stands for the total cell mass summing up the density of all of its progenies. **(c)** Sankey diagram showing the proportion of density outflowing from MEPs to downstream cell types or staying in the MEPs. The elongating bars of MEPs represents the expansion of MEP population due to self-proliferation. **(d)** Velocity stream plot showing the estimated differentiation rate as the velocity. To show the change of velocity field on the entire landscape over time, the diffusion map coordinates of all cells are taken by the drift module *v*(*s, t*) with t correspond to real time. **(e)** Boxplot of cellrank’s fate probability derived from velocity kernel. The shown fate probability was from MEPs towards terminal state Ery (brown) and terminal state Meg (blue). **(f)** Differential expression analysis comparing the early phase MEPs (from day3 to day12) with the late phase MEPs (day 49 and day 76). Genes with q-value higher than 0.05 and fold change greater than 2 (or smaller than -2) are considered as significant DEGs. Up-regulated DEGs are colored in pink and down-regulated DEGs in blue. **(g)** The low-output gene signature scores of MEPs at different timepoints.

Here, we focused on using CDT to study the progressive transition from an early megakaryocyte-biased progenitors to the balanced homeostatic state, as captured in our data. For simplicity, we focus on the bifurcation between the megakaryocyte and erythroid fates, yielding a reduced dataset comprising four populations: HSCs, Meg–Ery progenitors (MEPs), megakaryocyte progenitors, and erythrocyte progenitors. We reprocessed this subset and retrained the model on data spanning Day 3 to Day 76, thus capturing the time window post transition to the homeostatic state. Based on this setup, we performed density transport analysis for every MEP, a cell population characterised by Meg plus Ery bipotency. Density transport analysis of individual MEPs revealed dynamic shifts in progeny distribution over time (Fig. 6b). We annotated the intermediate cell states by their closest cells in the landscape and then aggregated the progeny density by cell type. In this way, we can quantify the mass of cell flux from MEP to erythrocytes, megakaryocytes, and self-renewing MEP compartments (Fig 6c). In the first two-time course, MEPs are generally less active in redistributing the cell mass, with a large fraction of descendants staying within MEPs. Between Day 12 and Day 27, MEPs acquired a transient megakaryocyte bias, which subsequently gave way to a balanced erythrocyte–megakaryocyte output from Day 49 onwards. Such transition in cell fate preference is reflected in the time-dependent differentiation rate, which essentially represents a time-evolving velocity field positioned in the low-dimensional space (Fig. 6d & Fig. S. 10a). To quantify the directionality implied by the velocity field, we calculated the fate probability of each cell in the landscape by using cellrank’s velocity kernel with the estimated differentiation rate as input.The resulting fate probability of MEPs shows a predominant fate bias toward megakaryocytes at the first three time points, increasing fate transition towards the erythroid lineage, ultimately resulting in a balanced lineage output after Day 49 (Fig. 6e).

At the molecular level, MEPs showed a time-evolving transcriptional signature that aligns with the shift in their function, as revealed by our CDT analysis. Differential expression analysis between MEPs from early-stage MEPs (Day 3-12) and late-stage MEPs (Day 49-76) uncovered a set of genes that are with the differentiation of multiple lineages. These included the marker genes for myeloid differentiation, such as Elane, Il 11 and Mpo (Rydzynska et al., 2023; Quesniaux et al., 1992; Nauseef et al., 1988), the lymphoid differentiation marker Cd52 (Fuhr et al., 2022; Dearden and Matutes, 2006) and Ctsg that involves in monocyte maturation (Cheung et al., 2021). Notably, within down-regulated genes, we found Slfn2 and Mecom, which serve to maintain HSPCs in a quiescent state (Warsi et al., 2016; Katsoulidis et al., 2009; Warsi et al., 2022; Ditadi and Sankaran, 2022). To move beyond considering lineage marker genes, we finally turned to the low output and multi-lineage signatures defined from in vivo analysis of barcoded HSCs Rodriguez-Fraticelli et al. (2020), which showed that the low output signature was higher during the early timepoints (Fig. 6g), consistent with our analysis that early cells is slower in accumulating density. By contrast, the multi-lineage signature (Fig. S. 10) was higher during later time points, where our analysis shows balanced lineage output. Taken together, this analysis provides substantive molecular evidence to corroborate our cellular flux model approach, specifically highlighting how early changes in MK production are accompanied by dynamic changes in underlying molecular programmes.

## Discussion

In this study, we introduce *pseudodynamics+* providing a true step-change in our ability to capture population dynamics for complex multi-lineage differentiation landscapes at single cell resolution. Bringing together single cell genomics with population scale cell-flux modelling substantially enhances the biological insights that can be gained from either approach alone. Cell flux modelling is commonly performed at the population level, thus lacking the much-enhanced granularity of single cell genomics. Moreover, input data are commonly devoid of deep molecular information (e.g. simple flow cytometry), which prohibits any connection of population-scale behaviours (growth, drift etc) with the underlying molecular machinery. Single cell genomics, on the other hand, commonly reports static snapshots, and even when time-series are generated, they lack information on population-scale flux dynamics. Importantly, it is is population-scale flux dynamics that are (i) perturbed in many disease settings, and (ii) need to be rectified to treat the disease. Bringing together single cell genomics with flux modelling of complex systems thus has the potential to provide both a single cell resolution and quantitative encapsulation of normal cell population-scale behaviour, as well as unlocking new avenues for therapeutic intervention.

We achieved the modelling population/tissue size dynamics over time and at single cell resolution by formulating a dynamic process of single-cell density change, which is governed by a partial differential equation. To solve this PDE, Pseudodynamics+ leverages the concept of physics-informed neural networks and parameterizes the rate of growth, drift, and diffusion with neural networks. The fitted behaviour networks then provide dynamic parameters across a single cell differentiation landscape for any arbitrary time point, which empowers (i) the interpolation of density landscapes and population/tissue sizes at unseen time points, (ii) simulation of cellular trajectories in a velocity field, and (iii) reconstruction of the continuous density transportation for any given cell along its differentiation trajectory. Finally, a bespoke drift-association test identifies genes whose onset or shutdown is associated with the most profound transcriptional changes during differentiation, thus devising a method for gene programme discovery that leverages previously unexplored population/tissue scale parameters. Pseudodynamics+ therefore constitutes an innovative and powerful framework that delivers multi-scale and actionable insights all the way from tissue to the molecular level.

When applied the development of embryonic thymocytes, *pseudodynamics+* estimates a high growth rate for DP cells that disagrees with pseudodynamics-v1 but better matches the cell cycle phase (Fig. 2e). Of note, the original pseudodynamics-v1 was built on a linear cell state coordinate (pseudotime), enforcing a zero drift term for the last pseudotime bin, essentially therefore enforcing a closed system. Instead, our method only builds a soft boundary by applying a loss term that penalises the mass flowing out of the system. Although the optimal strength of the boundary constraint can only be determined retrospectively, a soft boundary coupled with the open system assumption is intuitively more suitable for the thymocyte development data, because (i) the DP-cells will further differentiate to the single-positive stage and eventually exit from the thymus to other organs (Park et al., 2020a) , and (ii) our unobserved cells just beyond the measured trajectory can be assigned a smoothed value rather than be forced to immediately go down to zero drift.

Application of *pseudodynamics+* to the *in vivo* haematopoiesis data not only demonstrated how this approach can capture the flux parameters of complex multi-lineage differentiation processes, but also showed – perhaps unsurprisingly – that for long *in vivo* time courses of, in this case, 9 months, it is unlikely that models with fixed parameters will provide a good fit to the data (Fig S. 6). A particularly large degree of variation in the dynamic parameters was observed during the first couple of weeks of the time course. Two intuitive explanations come to mind: (i) a systemic perturbation by the tamoxifen treatment used to induce the Cre-dependent expression of the stable fluorescent marker, and (ii) heterogeneity in the HSC pool with HSC subtypes being distinguished by different kinetics. Tamoxifen treatment has previously been suggested to inhibit JAK-STAT signalling (Sánchez-Aguilera et al., 2014), and we previously reported a >50%-fold expansion of overall MEP and MkP-like progenitors Kucinski et al. (2024), thus matching cell types with the most elevated early growth rates (Fig 4).

The bone marrow HSC pool consists of functionally diverse HSC clones (Laurenti and Göttgens, 2018; Haas et al., 2018; Rodríguez-Correa et al., 2025). A recent HSC transplant study reported that fast-engrafting HSC clones displayed a megakaryocyte- and erythroid-bias compared with myeloid and lymphoid-biased slow-engrafting HSCs. Similarly, a non-canonical platelet differentiation pathway was recently reported whereby a subgroup of vWF-expressing HSCs was specialised to rapidly produce megakaryocytes and platelets under stress conditions (Sanjuan-Pla et al., 2013; Haas et al., 2015; Carrelha et al., 2024). Initially faster differentiation rates towards megakaryocytes in our dataset may reflect stem cell heterogeneity during homeostasis, where a small subset of rapidly differentiating megakaryocyte-biased HSCs dominate early cellular output, followed by a delayed contribution from the slower differentiating unbiased HSCs. Of note, our bespoke continuous density transport analysis identified distinct behaviours within the MEP progenitor pool (Fig. 6b), with megakaryocyte biassed behaviour followed by more balanced differentiation at later time points (Fig S. 10b). Together with the compositional change of vWF+ HSC and vWF+ progenitor cells (Fig. 5e), a model emerges whereby a temporal shift of HSC subtypes may propagate downstream to progenitors. Dissecting the precise impacts of tamoxifen treatment versus HSC heterogeneity will require the development of alternative experimental approaches.

*Pseudodynamics+* uses diffusion map coordinates rather than directly utilising gene expression as input due to the sparsity and difficulty of density estimation for high-dimensional data (Burkhardt et al., 2021; Otto et al., 2024). The lack of explicit mapping from cellular expression to cell dynamic rates makes it difficult, therefore, to directly quantify the contribution of each gene to cell flux parameters. Reversible dimensionality reduction Sha et al. (2024) has been applied to overcome this limitation, but the gradient-based feature attribution is rather sensitive to noise Sundararajan et al. (2017); Yang et al. (2023) and requires other feature selection techniques Huang et al. (2023) to infer candidate regulatory genes. Importantly, by linking molecular gene expression data with population-scale cell flux behaviours, *pseudodynamics+* nonetheless unlocks novel ways of identifying differentiation involving genes, as demonstrated here by correlating dynamic changes in differentiation speed (drift) with gene expression. As with other pipelines for defining contributing genes, our findings represent associations which may or may not have regulatory causality.

Taken together, *pseudodynamics+* bridges the gap between static single-cell snapshots and the dynamic, population-aware processes that govern tissue development and homeostasis, enabling insights into proliferation, differentiation, and lineage bias that were previously inaccessible from single-cell data alone. Beyond haematopoiesis, Pseudodynamics+ provides a generalizable approach for studying dynamic systems where both state transitions and population flux matter, offering a foundation for predictive modelling in regenerative biology, disease progression, and therapeutic intervention.

## Method

### *pseudodynamics+* models

In this study, we introduce *pseudodynamics+* to model the density shift of the single-cell population over time [1, 2, … , *T*] and the cell state space **S**. We improve the resolution of cell population dynamics to the cell state level and use cellular density *u*(*s, t*) to describe a continuous and infinitely dividable cell mass for the cell state s at time point t. The density function is scaled such that its integral over the state space **S** corresponds to the total population size at each time point, denoted as *N*_*t*_ = ⎰_*S*_ *u*(*s, t*)*ds*. This evolution in density, influenced by differentiation and growth, is described by an advection-reaction diffusion partial differential equation, as outlined in the equation. The density change driven by differentiation and growth is governed by an advection-reaction diffusion partial differential equation, which was introduced in the original *pseudodyanmics* with a dimension-less formulation of the PDE. Our work advances this framework to accommodate a multidimensional context, resulting in the system’s progression as follows:

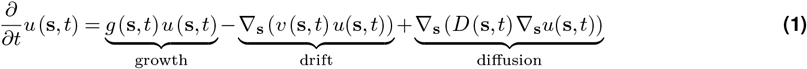

Here, the cell state **s** is a vector with *d* dimension. The time derivative of density is controlled by three different terms, including the net proliferation of the cell population with the growth rate *g*(*s, t*), the directed transition of cell state (drift) regulated by the differentiation rate *v*(*s, t*), and the stochastic redistribution of the density (diffusion) regulated by the diffusion rate *D*(*s, t*). With time series single-cell snapshots and population size, we can measure the density at the time of experiment. Given the disconnected density observation, *pseudodynamics+* solves an inverse problem of estimating the parameter *g*, *v* and *D* as functions of s and t, which we refer to as behaviour functions. Because of the complexity of behaviour function, we approximate the behaviour functions with three Multi-Layer Perceptrons (MLPs) *g*_*w*_(*s, t*) :*R* ^*d*^ ×[1, *T*]→*R*, *v*_*w*_(*s, t*) :*R* ^*d*^ ×[1, *T*]→*R* ^*d*^ and *D*_*w*_(*s, t*) :*R* ^*d*^ ×[1, *T*]→*R* ^*d*^, parameterized by *w*_*g*_, *w*_*v*_ and *w*_*D*_ respectively. The inverse problem thus turns into the estimation of the parameters of the behaviour function *w*_*{·}*_. A physics-informed neural network, a novel deep-learning based method to solve complex PDE problem, is employed to solve the inverse problem. A surrogate network *u*_*θ*_(*s, t*) :*d*×[1, *T*]→*R* is introduced accordingly to approximate the actual density function *u*_*θ*_(*s, t*)*≈ u*(*s, t*), is another MLP taking in the cell state *s* and the time *t* to predict the density. By minimizing the following forward prediction loss *L*_*θ*_ (Eq.2), the surrogate density network can learn to match the observed density.

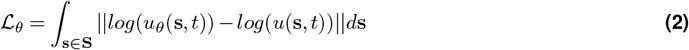

To ensure that the predicted density at the observed time aligns with the dynamics that obey the physics law defined in Eq.1, we apply a residual loss *L*_*r*_ (Eq.4) to constrain the surrogate network.

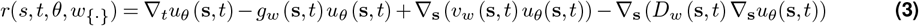

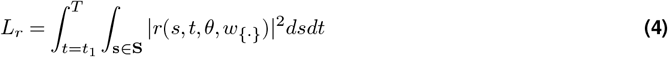

Eq.3 defines the residue term *r* of the surrogate network, which shares the parameters *θ* and *w*_*{·}*_. The residue term measures where and when a cell violates the PDE system. Optimising the residue across the cell space S and at any time point *t* provides smooth density interpolation between the observed time point.

### integration with NeuralODE

The density change between two time points can be simulated using NeuralODE Chen et al. (2018b) over the right hand side of Eq.1, which can be decoupled into surrogate network dependent terms and behavior network dependent terms Eq.5

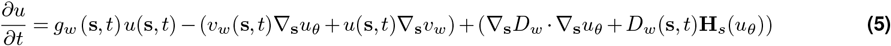

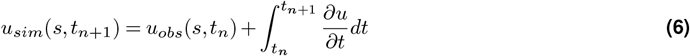

where ∇ _*s*_*u*_*θ*_ is the first order derivative and **H** _*s*_(*u*_*θ*_) =∇ _*s*_(∇_*s*_ *u*_*θ*_) is the second order derivative of the surrogate network relying on the auto-grad calculation enabled by modern deep learning platform Paszke et al. (2017). Similarly, we can approximate the cell state gradient of the behaviour network ∇ _*s*_*v*_*w*_ and ∇ _*s*_ *D*_*w*_ using autograd. A simulation loss *L*_*sim*_ is calculated to quantify the density error accumulated on the integral due to the estimated dynamic parameters:

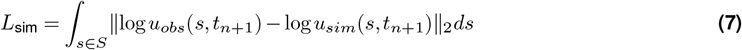

where *L*_*sim*_ is defined by the norm-2 error between the simulated density *u*_*sim*_(*s, t*_*n*+1_)and the observed density *u*_*obs*_(*s, t*_*n*+1_) at the next time point *t*_*n*+1_ on the logarithmic scale (Eq.7), with which the behaviour functions are tuned more precisely to align with the data set.

### Adapting *pseudodynamics+* for single-cell data

*Pseudodynamic+* constructs its total loss *L*_*total*_ as a weighted sum of several terms of constitution loss, each encoding different physical or statistical constraints to regularise and inform the inference of PDE parameters at the population level. We defined the total loss as follows:

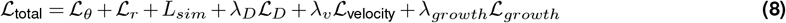

where each *λ* is the hyperparameter to weight the corresponding loss term. Apart from the PINN-loss (*L*_*r*_) and the density fitting loss (*L*_*θ*_, *L*_sim_), we introduced several regularisation terms to further constrain the estimated behaviour to fit the single-cell data. We discretised the cell state space **S** by taking the observed cells as the whole set. Thereby summing up the unnormlised density across the single-cell dataset with *n* cells equals to the total population size *N*_*t*_ = ∑_*s∈***S**_ *u*(*s, t*). Specifically, ℒ_*D*_ is to restrict the value of the diffusion so that the density change can be attributed more to the growth and drift terms. The velocity loss ℒ _*v*_ (Eq. 10) is to constrain the direction of *v* to comply with the local geometry so that simulated cell state can stay in the data:

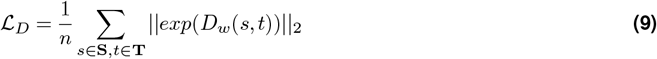

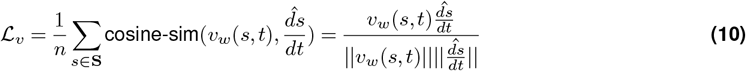

where 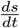 is the local cell state change inferred from the single cell data set. RNA-velocity is taken when it is available; otherwise we sample the cell state from the K nearest neighbours according to their distance to the user-provided root cell. With this regularisation, our framework can more focus on estimating the vector sum, which is a scaler showing the differentiation rate.

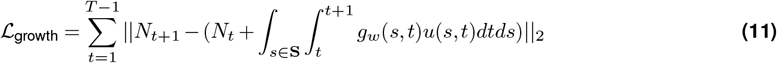

The growth loss ℒ_growth_ to control how well the growth-driven cell mass gain can match the measurement. Hypothetically, when the system is closed, all density gain over time is only contributed by growth, while drift and diffusion terms only redistribute the density across different parts within the landscape. The hyperparameter *λ*_growth_ = 0 stands for an open system, where the outflux to the unobserved cell state is allowed.

### Data preprocessing

After quality control and batch corretion, the diffusion map space is calculated using `palantir.utils.run_diffusion_maps`, which gives rise to the cell state space **S** our model depends on. For each observed time point *t* and cells **S** _*t*_, we use the Gaussian kernel density estimator (kde) of the package `scipy` to obtain the density function *kde*_*t*_ (*s*). The raw density over the space is obtained by applying a time-specific kde function to the whole space. The raw density values are later rescaled so that their sum matches the experimentally observed or desired total population size for that time point.

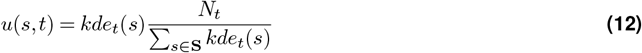

### Model training

The cell state space **S** is randomly split into training, validation, and test set at a default ratio of (0.8,0.1,0.1). When a specific time point *t* is held out, the cell states *s*_*t*_ from the masked time point are placed in the test set and the density *u*(*s, t*) is held out for the model during training. Without prior knowledge of the single cell dataset, the default hyperparameters are as follows: *λ*_*D*_ = 1, *λ*_*v*_ = 0.01, and *λ*_growth_ = 0. The parameters of the surrogate network *θ* and the behaviour networks **w**_*{·}*_ were optimized by `Adam` optimizer with the default learning rate of 3*e* − 4. `Dopri5` solver is used by default for the NeuralODE integral with a default tolerence value of 1*e*−4.

### Cell state transition simulation via velocity-field

Cell state transitions are simulated by integrating the learned velocity field **v** _**w**_(**s**, *t*) over continuous time, where *s* represents the *d*-dimensional diffusion map space and *t* denotes experimental time. Two simulation paradigms are implemented, with different biological assumptions about the cellular differentiation process.

In the deterministic setting, cell state trajectories follow the ordinary differential equation (ODE):

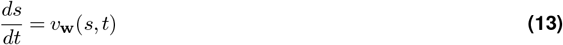

where **v**_**w**_ (**s**, *t*) is the velocity field learned from the physics-informed neural network (PINN) framework. This formulation assumes that cellular transitions are governed entirely by deterministic forces encoded in the velocity field.

For the advanced stochastic formulation, cell state dynamics incorporate both directed velocity and random diffusion processes:

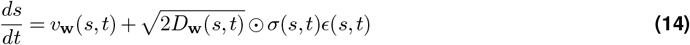

where **D**(**s**, *t*) is the learned diffusion tensor from the PINN model; *σ*(**s**, *t*) represents a user-defined noise schedule

function (default: constant unity), *ϵ*(*t*)∼ 𝒩(**0**,**I**) is multivariate white noise by default. The integration employs the adaptive ODE solvers `Dopri5` from the `torchdiffeq` library with a tolerant of 1*e*−4 by default. When encountering numeric underflow, we switched to fixed-step solver `Rk4` with 100 steps over the integration time.

### Cell type labelling for simulated trajectory

Let [*s*_1_, *s*_2_, *· · ·, s*_*n*_] denote the vector of simulated cell states. For cell type assignment, a supervised nearest-neighbor is applied to transfer the cell type label from the reference dataset. For every simulated state *s*_*i*_ in the trajectory, the algorithm queries the nearest cell from the reference dataset and transfer their labels using `annoy`, a fast approximate nearest-neighbour searching method. The indices corresponding to the nearest reference cells to the query states are retrieved using the Euclidean in diffusion map coordinates. The assigned cell type for simulated state is determined by majority vote among the cell type labels of the nearest reference neighbors.

### Evalute simulated trajecoty in Weinreb data

The clonal annotation in data by Weinreb et al., the lineage-traced hematopoiesis single cell data, recovers the history where cells differentiate to at the next time point, which serves as a basis for evaluating our simulated trajectory. Fate bias quantifies the probability that a cell, simulated from an initial state or ensemble, commits to a particular fate (cell type) at its terminal state. The cell fate of the cell is defined by the most frequent cell type of the clone it belongs to. Here we simulated trajectories for every cell at day 2 and finished at day 6, and assigned a cell type label to the last simulated state. The fate prediction accuracy is calculated between the assigned cell type and the true cell type using `sklearn.metrics.accuracy_score`. The W-2 distance was calculated between the actual cell state at Day 6 and the trajectory simulated over a time span from day 4 to 6. The simulation starting points are set to cells at day 4 from clone 3869, 2673, and 2831, which are held out from training. The W-2 distance is implemented using `pot.sinkhorn2` with settings `power=2` and `reg=1`.

### Drift-association test

#### Project the estimated parameters on the pseudo-time axis

The model-inferred growth rate *g* and vector field *v* sits in the high-dimensional cell representation space. To analyse how cellular dynamics evolve along developmental trajectories, the parameters can be projected onto a univariate pseudotime axis. Each cell was assigned to a lineage according to the fate probability defined from package `palantir` or `cellrank` following Kucinski et al. (2024). For each lineage, cells belonging to that lineage were subsetted, giving rise to the lineage specific cell state vector. Their pseudotime values, obtained from either `scanpy` or `palantir`, were binned into 100 uniform intervals. When given a timepoint for time-dependent models, the growth rate and the differentiation rate are estimated using the cell state vector. For differentiation rate specifically, the vector sum is calculated to convert a multi-dimensional vector field into a univariate scaler representing the speed along the trajectory. The average value of the parameter for each bin was computed, resulting in a smoothed trajectory of the dynamical parameter over pseudotime. The binned pseudotime and binned parameters were taken as anchors to perform linear interpolation of each single-cell’s projected parameter based on the actual pseudotime value.

#### Identification of the pseudotime window with the steepest velocity

To perform association test, we first computed the time derivative of the projected differentiation rate 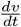 to identify the time window with largest slope. For each lineage and timepoint, we defined the projected velocity profile **v**(*t*) and computed its numerical derivative with respect to pseudotime via finite differences:

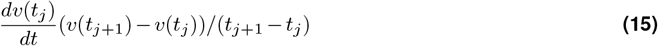

To identify regions of maximum dynamical change, a window of length *h* (by default 30 pseudotime bins) is slid over the pseudotime axis. For each window, calculated the mean absolute slope and selected the window with the maximal value, returning pseudotime interval 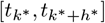 as the region of steepest velocity. Fitting Generalized Additive Model (GAM) and Trajectory differential expression test Next, we computed the smoothed gene expression over the trajectory of the selected lineage, to comply with the projected differentiation rate. We modelled the relationship between gene expression and pseudotime using a Generalized Additive Model (GAM) framework similar to `tradeseq`. For gene expression matrix *X* ∈ ℝ ^*N*×*G*^ , we fit a GAM per gene using its observed expression and cell pseudotime. The model for gene *g* follows the negative bionomical (NB)

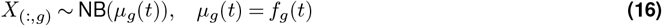

where the mean *µ*_*g*_ is a spline function *f*_*g*_(*t*) of pseudotime *t*. To better integrate with our modelling framework, we basically reimplemented the `tradseq` GAM fitting in python via the `statsmodels` library. *f*_*g*_ is realised by a cubic B-spline basis (`statsmodels.gam.api.BSplines` with `n_knots=7` by default). Restricting to the previously identified pseudotime window 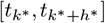,we performed the wald-test using the fitted GAM like `tradeseq` Van den Berge et al. (2020). For each gene, the contrast matrix *L* was defined as the differences of the NB mean. The wald test statistic *Wald* was calculated following:

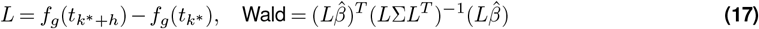

where *β* represents the coefficients of the GAM and Σ is the covariance matrix. The p-value was retrieved referring to the Chi-square distribution with python function `scipy.stats.chi2`. FDR-value was obtained using the `multipletests` function from `statsmodels` library and a threshold of `FDR<=0.01` was applied to claim significance.

### Quantification of Gene-Drift association

As **v** is considered the time derivative of cell state transition, 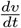 is therefore the second order derivative of cell state over time, which we hypothesised was proportional to the force driving cell state transition. To identify molecular correlates of this force term, we quantified associations between gene expression and 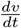. Within significantly differentially expressed genes, the linear association between the smoothed expression (from GAM) and 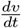 was captured with Pearson correlation coefficient. The non-linear association was measured using mutual information (MI) from `sklearn.feature_selection.mutual_info_regression`. Finally, we combined the trajectory DEG test and association to filter genes that are associated with the model-estimated differentiation rate. For transcription factors, selection criteria required a mean log fold-change >1, FDR<=0.01 and MI > 0.7. For all other genes, the criteria for log-foldchange was increased to 2. In the figures, genes shown were ranked by their MI from high to low.

### Continuous Density Transport Analysis

In this study, we developed continuous density transport (CDT) that relied on multiple dynamics parameters to characterise density redistribution from one cell to all its progeny. CDT distinguishes itself from dynamic optimal transport in that it is a post-training simulation for individual cell, while dynamic optimal transport is for model training on the entire dataset. Our CDT analysis couples individual cell state transitions with population-level density dynamics to capture both local cellular behaviours and global population flows. Given a starting cell *s*_0_ at time *t*_0_, its transition trajectory [*s*_1_, *s*_2_, *· · ·, s*_*n*_] was simulated throughout a time window [*t*_1_, *t*_2_, *· · ·, t*_*n*_] in the deterministic setting. CDT constructs a transport matrix **T**∈ *R* ^*n*×*n*^ for each cell, where T[j, i] represents the density of progeny cells in state *s*_*i*_ at time point *t*_*j*_. Along the trajectory constitutes the source cell state *s*_*i*_ and the target cell state *s*_*i*+1_ of every transport event. There are three components of the simulation, including the density outflow *u*_*i*→*i*+1,Δ*t*_ from current cell state to the next state during time interval Δ*t*=*t*_*j*+1_ −*t*_*j*_ =*t*_*j*_ −*t*_*j*−1_, *j > i*, the inflow receiving from the previous state is *u*_*i*−1→*i*,Δ*t*_ and the density retention *u*_*i*→*i*,Δ*t*_ representing the untransported cell mass. The density of progeny cells at state *s*_*i*_ originated from the starting cell *s*_0_ is as follows:

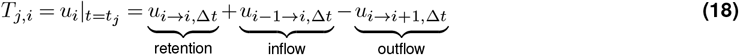

As a key assumption, the direction of the density transport is determined locally by individual cell alone yet the cell mass being transported also dependent on the global density difference between the source and target state. The momentum at cell state was computed via:

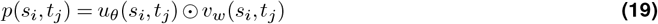

where *u*_*θ*_ is the trained density network that return the density at population wise and *v*_*w*_ is the fitted behaviour network. We estimated the global drift using finite difference approximation:

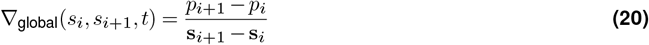

This global drift term approximates the gradient of momentum across the state space trajectory. The density outflow from source *i* to target *i* + 1 during interval [*t*_*j*_, *t*_*j*+1_]is:

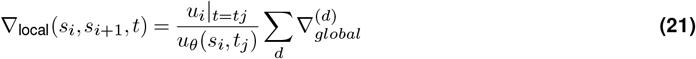

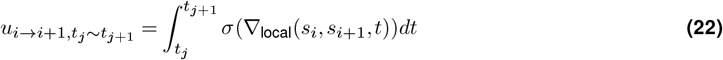

where *σ* is the ReLU activation function that ensures non-negative density flow. The density retained at state i is obtained by accounting for self-proliferation and outflow:

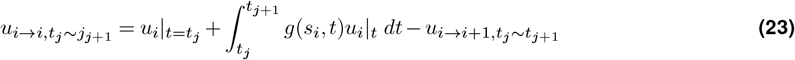

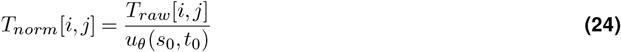

The construction of the transport matrix **T** follows an iterative procedure from *t*_1_ to *T*_*n*_. At intermediate timepoint *t*_*j*_, the density transport happened within cell states *s*_1_ to *s*_*j*_ within each iteration. Finally, the raw transport matrix is normalized to account for the difference of starting cell density.

## Code Availability

*psuedodynamics+* is implemented as a standard Python package, and the source codes are freely available at https://github.com/Gottgens-lab/pseudodynamics_plus. All analysis notebooks are also included in the repository for reproducibility.

## Author Contribution

B.G conceived the project. W.Z. designed and implemented the *pseudodyanmics+* model and the downstream applications. M.B helped with the mathmatic formulation of the model and the analysis. All the single-cell data analysis and interpretation was performed by W.Z under the supervision of F.T, N.W and B.G.

## Supplementary Figures

**Figure S. 1.**
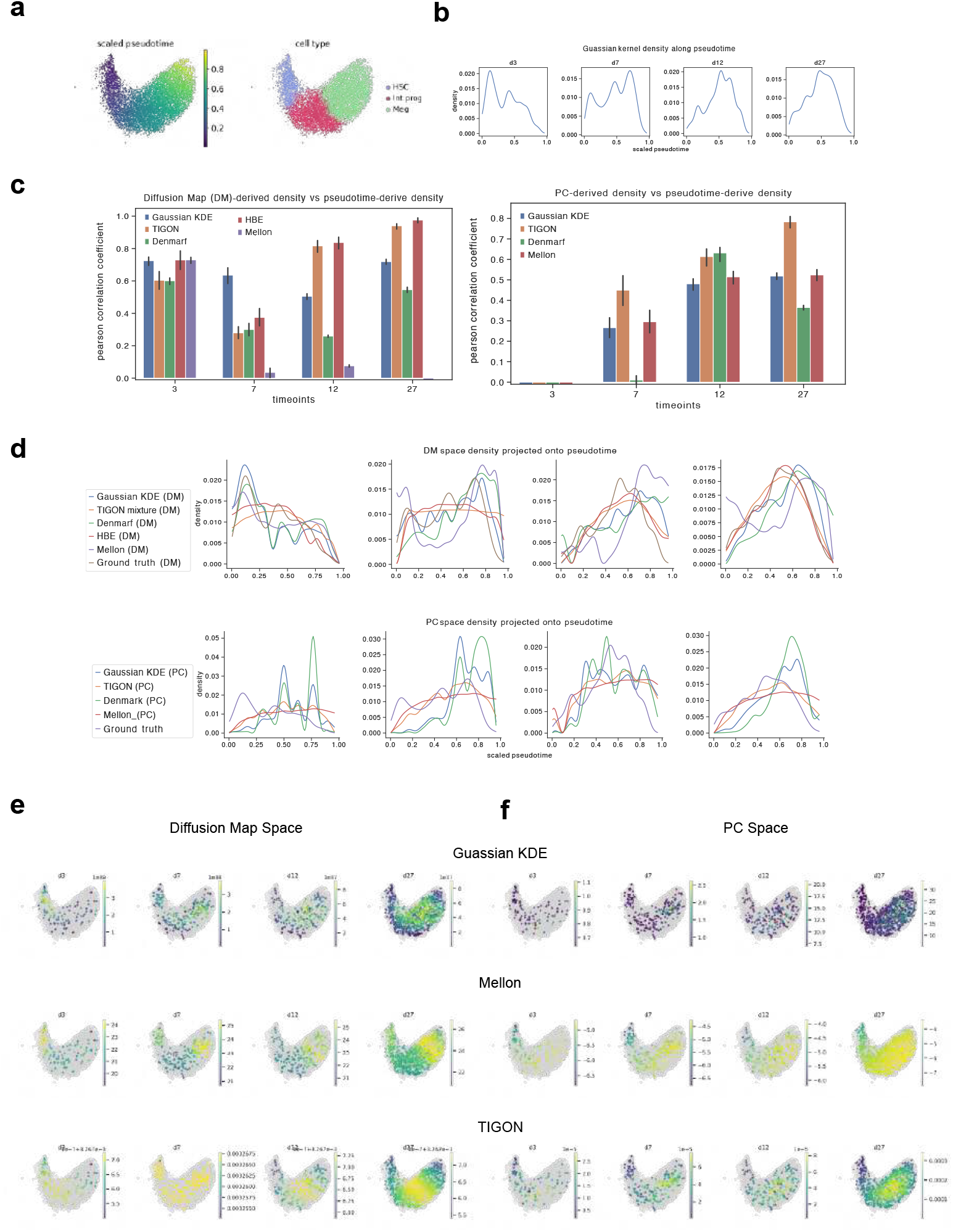
Benchmarking of different density estimation on different cell state space. In the megakaryocyte-lineage differentiation data, we benchmark the both the dimension reduction space and the density estimation function on their ability to recover a distribution shift along the trajectory over five time points. The UMAP recapitulates the megakaryocyte trajectory with stem cell on the very right end, progenitor cell in the middle and mature megakaryocte cells on the left. **(a)**. Umap visualizing a megakaryocte development dataset with progressing pseudotime and cell type over time. **(b)**. Line plot showing the gaussian kernel density estimation (KDE) on the space of 100 pseudotime bins, which each sub panel showing the density distribution of cell from different time points. **(c)**. Pearson correlation coefficient between the projected density from different methods and the pseudotime-derived density on different trails of bandwidth parameters. At each time point, the pseudotime density (b) is taken as the baseline against which the other methods and spaces are compared. On the left shows performance on density estimated from Diffusion map space, while that from PC space is show on the right. **(d)**. Line plot showing the density projected onto pseudotime bins. DM space density is placed on top and PC space density on the bottom. Sub-panels from left to right show result for D3, D7, D27 and D49 respectively. **(e-f)**. Density estimation results on multi-dimensional space. UMAP on the upper panel shows the density of every cells.

**Figure S. 2.**
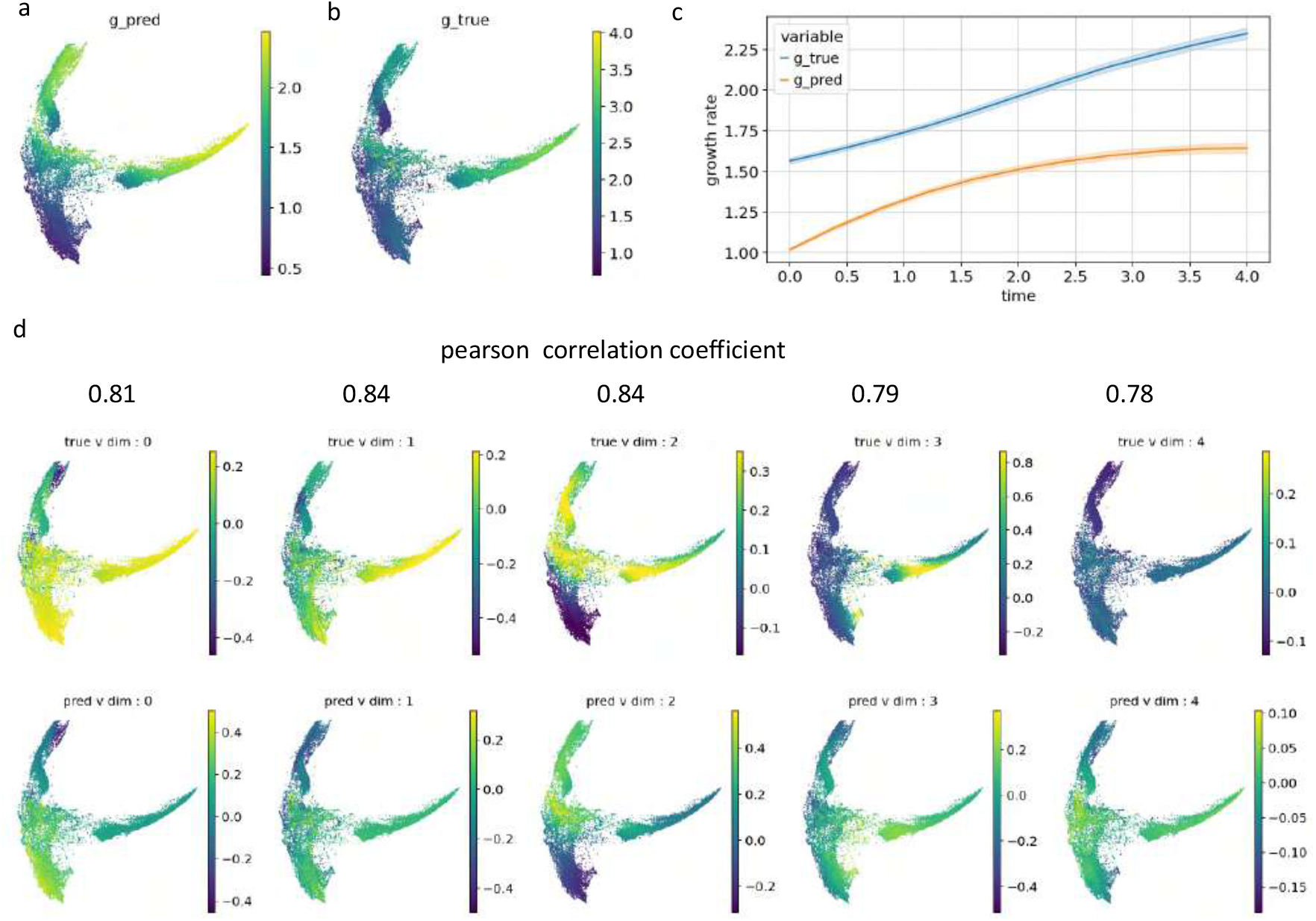
Evaluation of inferred growth parameter g and drift parameter v with the synthetic dataset. A synthetic dataset with known time- and state-dependent parameters was constructed. The dataset sits in a 5-dimensional space and the UMAP dimension reduction is performed for visualization. **(a-b)**. UMAP visualization of the ground true growth rate and the model prediction. **(c)**. Lineplot showing the average growth rate changes at time-wise. **(d)**. UMAP visualization the ground true and inferred differentiation rate per dimension. The Pearson Correlation Coefficients between the two rates per dimension are shown.

**Figure S. 3.**
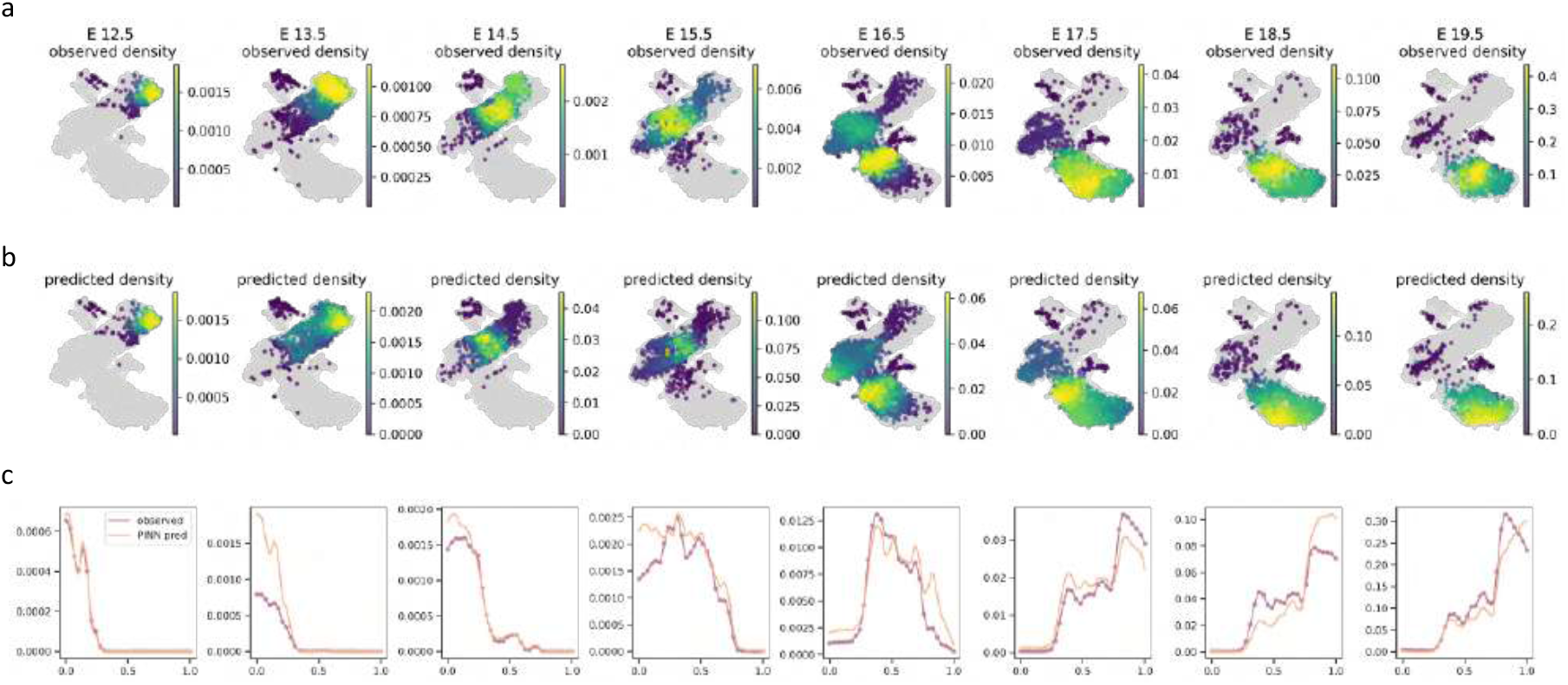
Evaluating the reconstructed density in mouse thymus dataset. **(a-b)**. UMAP visualization of the ground true cell density for each time points and the density simulated by *pseudodyanmics+* is shown at the bottom (b). **(c)**. Comparing the density fitting results along the scaled pseudotime axis.

**Figure S. 4.**
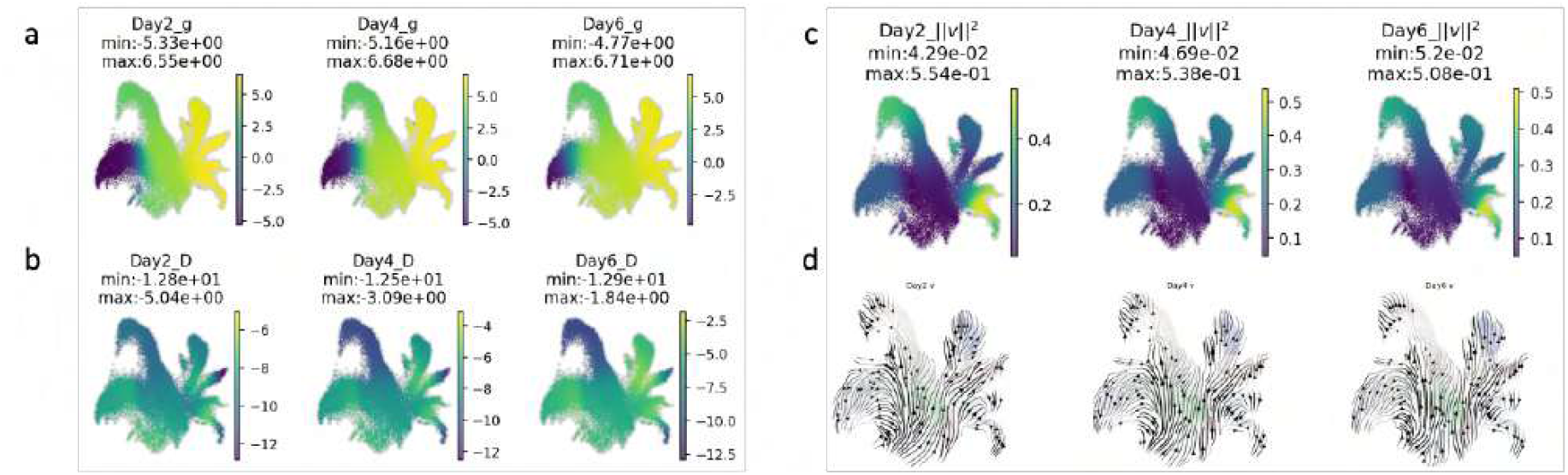
The dynamics parameters inferred by *pseudodynamics+* in Weinreb et al. data. **(a-b)**.UMAP visualization of the model inferred growth rate *g*(*s, t*)and diffusion rate *D*(*s, t*)with *S* using the entire cell state space and *t* is set to observed time points. **(c)**. Vector sum (L2-norm) of the differentiation rate *v*(*s, t*), which is original a 5-dimensinal vector per cell. Dynamic parameters are predicted across the whole landscape. **(d)**. The direction of drift is shown by velocity stream plot. Note that the model estimated differentiation rate is a multi-dimensional vector denoting the velocity of the cell in the diffusion map space.

**Figure S. 5.**
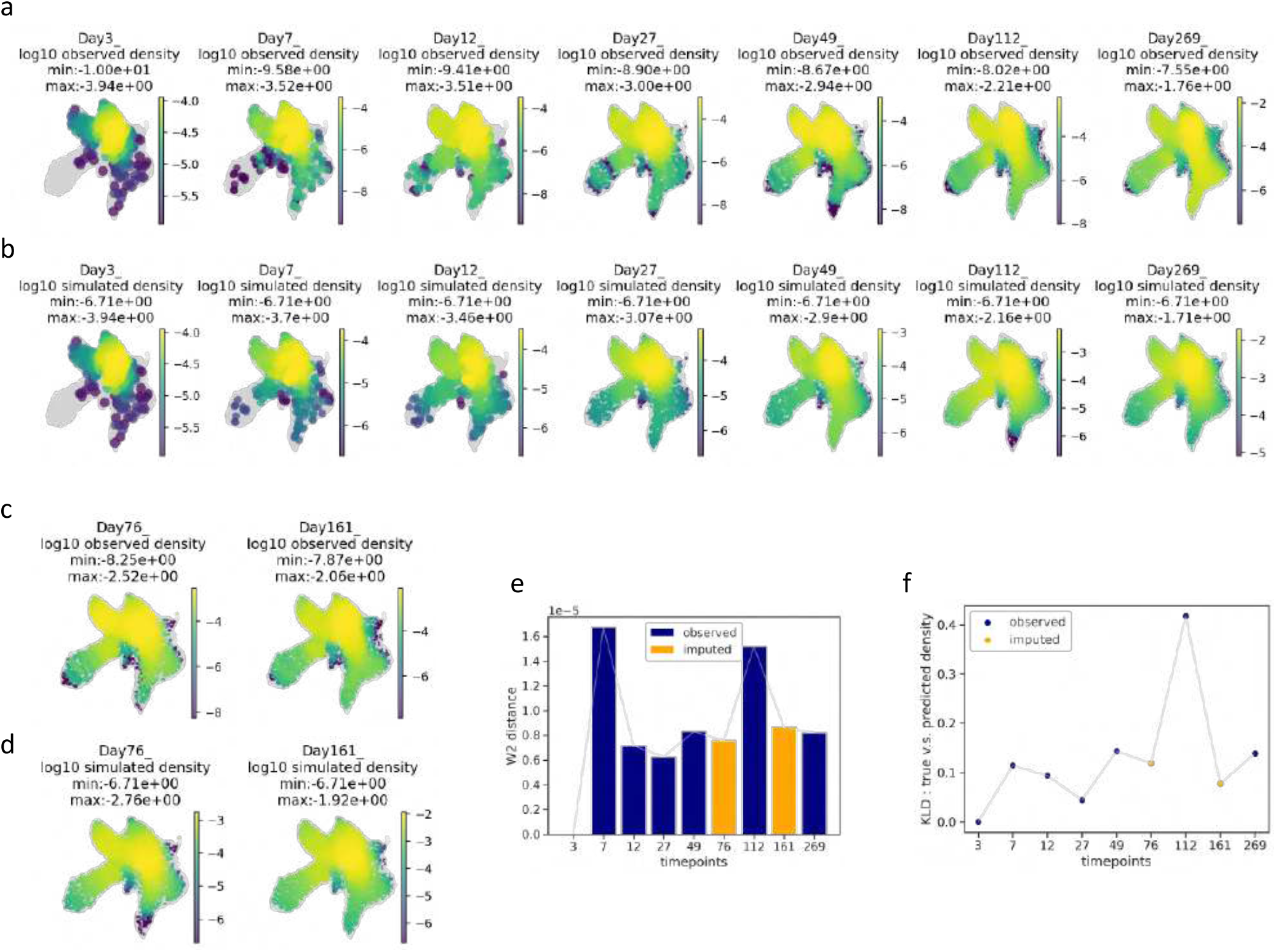
Evaluation of PINT Density Simulation Performance on in vivo Hematopoiesis Data. **(a, b)** UMAP visualizations of observed and predicted log10 densities for timepoints included in the training data. **(c, d)** UMAP visualizations of observed and predicted log10 densities for held-out timepoints. **(e, f)** Evaluation metrics comparing observed and predicted densities: (e) Wasserstein distance (W2) and (f) Kullback-Leibler divergence (KLD) across different timepoints.

**Figure S. 6.**
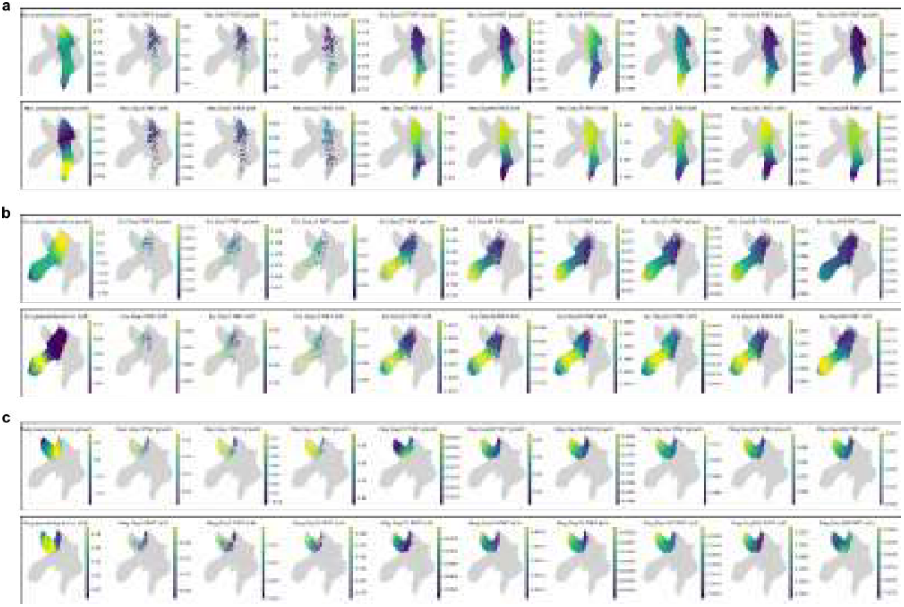
Comparing time-sensitive PINT derived dynamic paramters with pseudodynamics static parameters for Neu Ery and Meg lineage. The growth rate is placed as the upper row, and vector summed differentiation rate as at the lower row within each panel. Within each row, the leftmost column shows the parameters inferred by pseudodynamics (static model). The second column and onwards show PINT inferred parameters evaluated at each timepoint.

**Figure S. 7.**
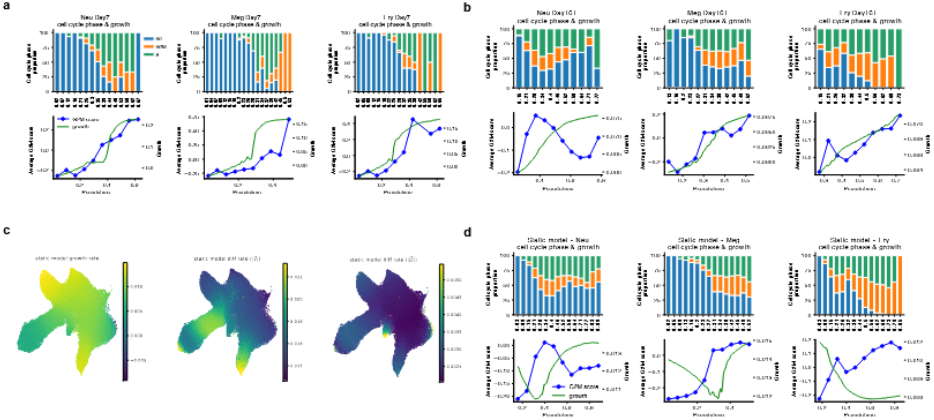
Evaluation of inferred dynamic parameters based on lineage-specific cell cycle analysis. **(a-b)** The stacked bar plots on the top show the compositional changes of cell cycle phase across pseudotime intervals. The lower line plots show the value of projected growth rate (y-axis on the right) as a function of pseudotime and average G2M score (y-axis on the left) over pseudotime intervals. Within each panel, columns from left to right stand for Neu , Meg and Ery lineage respectively. **(c)** The dynamic parameter estimated by our static model. **(d)** The cell cycle phase composition and G2M score along the trajectory. The growth rate of each lineage is coloured in green.

**Figure S. 8.**
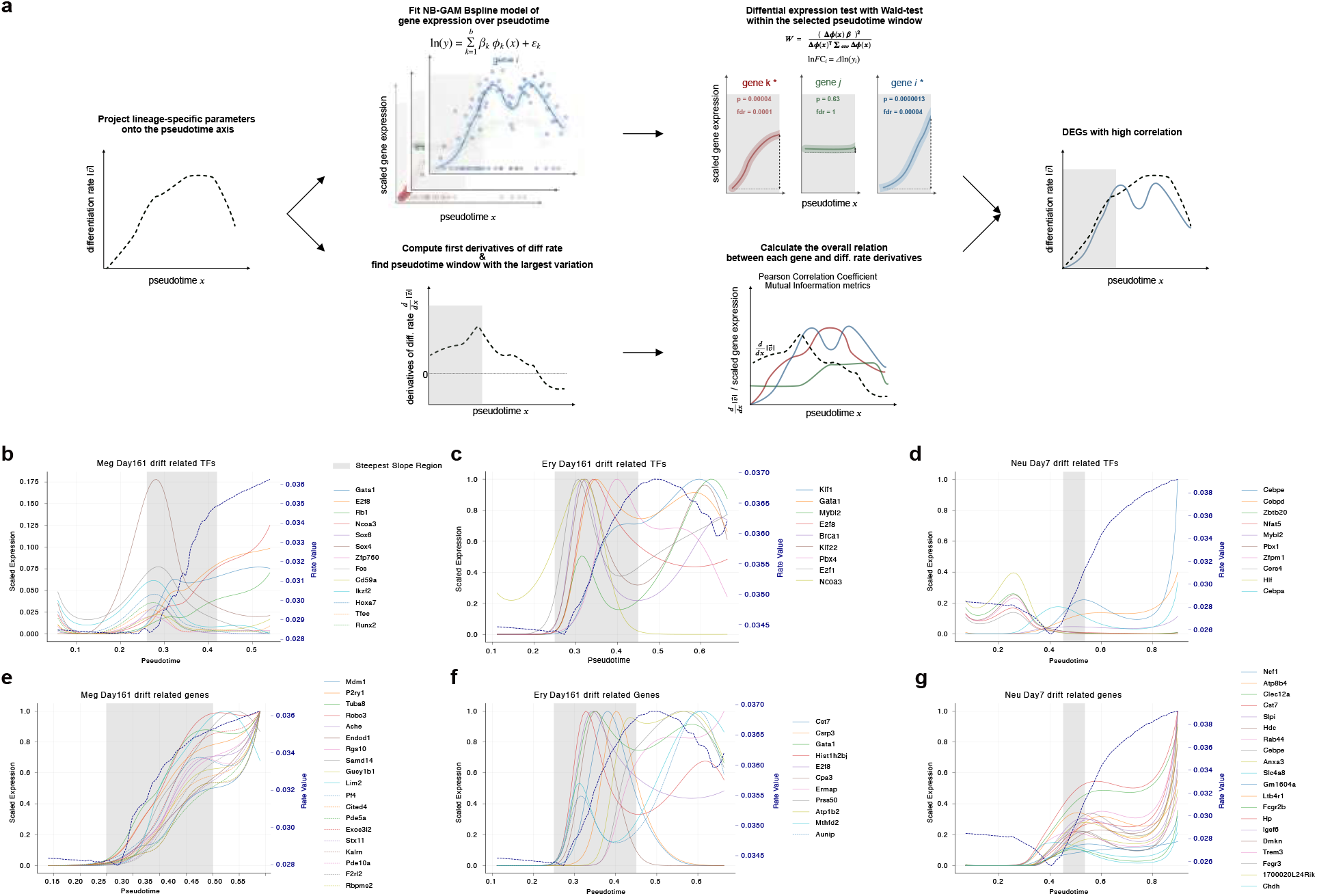
The drift associated genes for three main lineages of *in vivo* haematopoiesis data.. **(a)** The pipeline for searching and ranking associated genes when given the lineage-projected differentiation rate 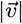. The workflow on the top shows the steps of fitting GAM gene curve and performing trajectory differential expression test within the given pseudotime interval. The workflow at the bottom shows the calculation of gradience of differentiation rate over pseudotime 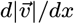 and the linear and non-linear correlation metrics between gene trend and the gradience 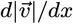. Genes were then filtered and ranked by combining the results from the two processes. **(b-g)** The curve plots showing the most associated TFs ( the upper row) and the genes (the lower row). From left to right displays the results for Meg, Ery and Neu lineages. The differentiation rate is shown as a bold dashed line with y-axis on the right and the other curves stand for the genes. The shaded area indicates the region where the diff. rate exhibits the steepest slope (the highest mean gradience).

**Figure S. 9.**
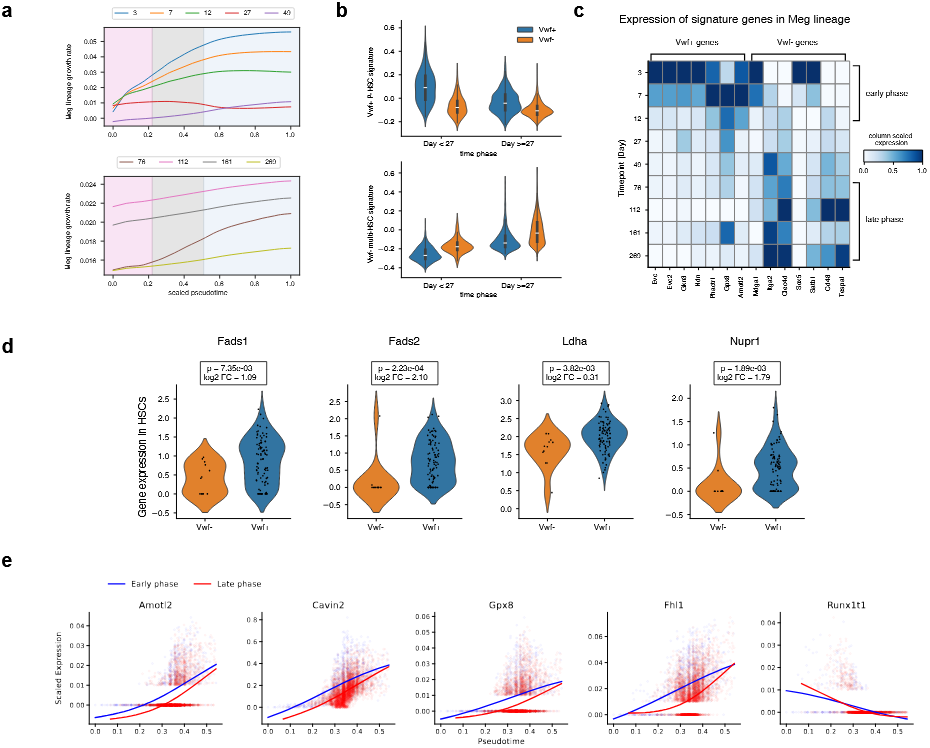
Vwf+ HSC population expression signature analyses.. **(a)** Growth curve of the megakaryocyte lineage plotted against pseudotime. **(b)** Violin plot showing the signature score of active genes in Vwf+ HSC transplantation and that in Vwf- multi-lineage transplantation. Only the score of HSC cells were displayed in the distribution and cells were grouped by whether collected before Day 27 or after Day27. **(c)** Heatmap showing the scaled expression of selected signature gene over time. The expression level was scaled for each gene across timepoints. Only the top 7 genes from each condition were shown. **(d)** The expression level of 4 mTol pathway related genes that were shortlisted as differentially expressed between the two condition. **(e)** Gene trend of differentially

**Figure S. 10.**
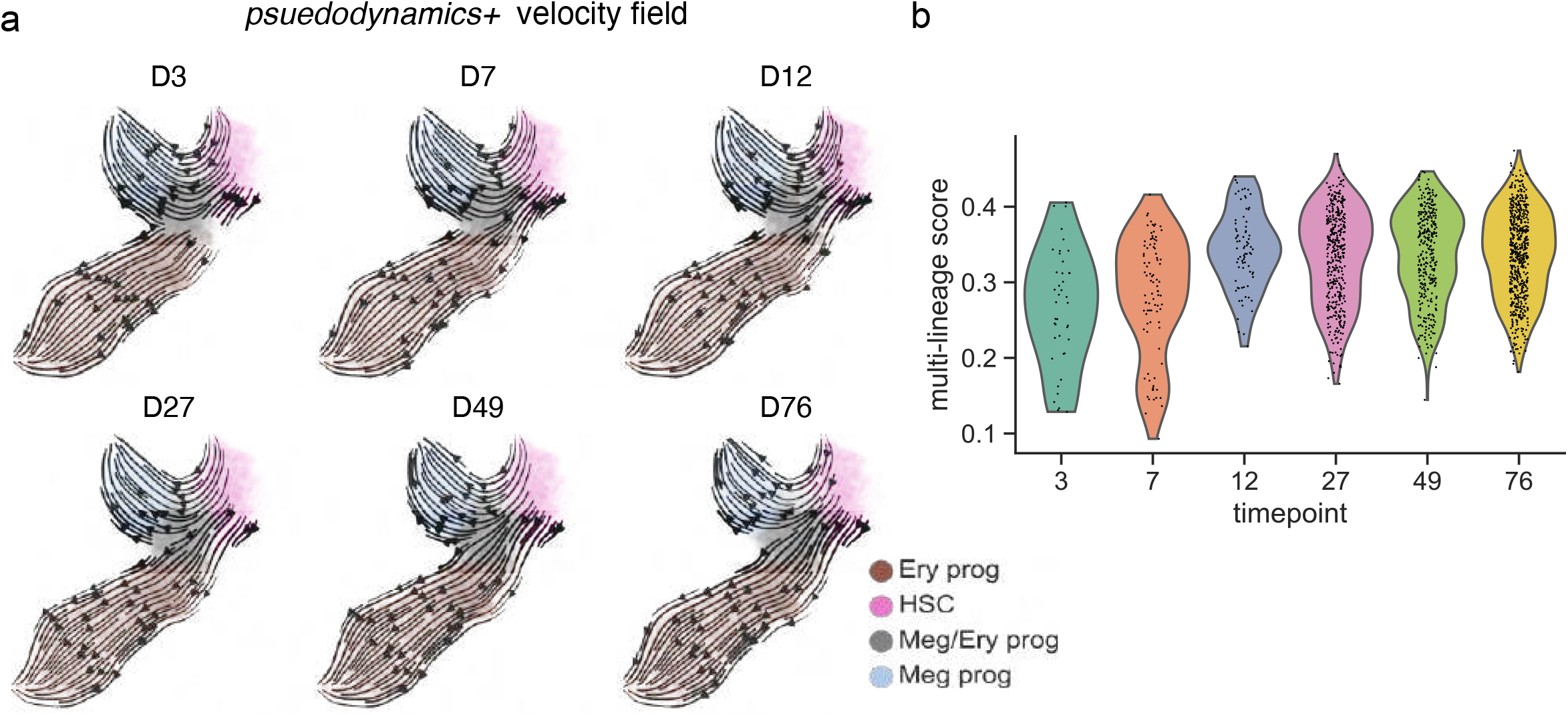
Temporal transition of MEPs in velocity field and lineage preference. **(a)** Visualization of pseudodynamics+ esitmated time dependent velocity field. From Day 3 to Day 76, the velocity arrows showed prominent changes in their direction around the MEPs area. **(b)** Violin plot showing the distribution of multi-lineage gene signature score for MEPs population over time points.

